# MAP7 family proteins regulate kinesin-1 recruitment and activation

**DOI:** 10.1101/388512

**Authors:** Peter Jan Hooikaas, Maud Martin, Tobias Mühlethaler, Gert-Jan Kuijntjes, Cathelijn A.E. Peeters, Eugene A. Katrukha, Luca Ferrari, Riccardo Stucchi, Daan G.F. Verhagen, Wilhelmina E. van Riel, Ilya Grigoriev, A.F. Maarten Altelaar, Casper C. Hoogenraad, Stefan G.D. Rüdiger, Michel O. Steinmetz, Lukas C. Kapitein, Anna Akhmanova

## Abstract

Kinesin-1 is responsible for microtubule-based transport of numerous cellular cargoes. Here, we explored the regulation of kinesin-1 by MAP7 proteins. We found that all four mammalian MAP7 family members bind to kinesin-1. In HeLa cells, MAP7, MAP7D1 and MAP7D3 act redundantly to enable kinesin-1-dependent transport and microtubule recruitment of the truncated kinesin-1 KIF5B-560, which contains the stalk but not the cargo-binding and autoregulatory regions. In vitro, purified MAP7 and MAP7D3 increase microtubule landing rate and processivity of kinesin-1 through transient association with the motor. MAP7 proteins promote binding of kinesin-1 to microtubules both directly, through the N-terminal microtubule-binding domain and unstructured linker region, and indirectly, through an allosteric effect exerted by the kinesin-binding C-terminal domain. Compared to MAP7, MAP7D3 has a higher affinity for kinesin-1 and a lower affinity for microtubules and, unlike MAP7, can be co-transported with the motor. We propose that MAP7 proteins are microtubule-tethered kinesin-1 activators, with which the motor transiently interacts as it moves along microtubules.

**Summary:** A combination of experiments in cells and in vitro reconstitution assays demonstrated that mammalian MAP7 family proteins act redundantly to activate kinesin-1 and promote its microtubule binding and processivity by transiently associating with the stalk region of the motor.

## Introduction

Kinesins are molecular motors responsible for the transport of different organelles and macromolecular complexes along microtubules (MTs) and for controlling MT organization and dynamics (Hirokawa and Tanaka, 2015; Verhey et al., 2011). The spatial and temporal control of kinesin localization and activity depends on numerous factors, such as cargo adaptors, post-translational modifications and the interactions with MT-associated proteins (MAPs) (Akhmanova and Hammer, 2010; Barlan and Gelfand, 2017; Fu and Holzbaur, 2014; Verhey and Hammond, 2009).

Kinesin-1 is the major MT plus-end directed motor involved in a broad variety of transport processes (Akhmanova and Hammer, 2010; Hirokawa and Tanaka, 2015; Verhey et al., 2011). This motor is well known to be regulated by different MAPs. Neuronal MAPs tau and MAP2 inhibit kinesin-1-driven motility (Dixit et al., 2008; Ebneth et al., 1998; Gumy et al., 2017; Monroy et al., 2018; Seitz et al., 2002; Trinczek et al., 1999; Vershinin et al., 2007). In contrast, MAP7 family members are firmly established to be positive regulators of kinesin-1 (Barlan et al., 2013; Metivier et al., 2018; Metzger et al., 2012; Monroy et al., 2018; Sung et al., 2008). MAP7 proteins are represented by a single homologue, ensconsin, in flies and by four isoforms encoded by different genes, MAP7, MAP7D1, MAP7D2 and MAP7D3, in mammals (Bulinski and Bossler, 1994; Metzger et al., 2012; Yadav et al., 2014). All MAP7 family members have a similar organization, with two conserved domains that are predicted to be helical, connected by an unstructured linker. The N-terminal domain of MAP7 proteins strongly interacts with MTs, while the C-terminal domain binds to the stalk region of kinesin-1 (Metzger et al., 2012; Monroy et al., 2018; Sun et al., 2011) (Fig. 1A). Additional regions with MT affinity were found in the linker of MAP7 and the C-terminal part of MAP7D3 (Tymanskyj et al., 2018; Yadav et al., 2014). In flies, ensconsin is an essential kinesin-1 cofactor required for numerous processes ranging from organelle transport to MT sliding (Barlan et al., 2013; Metivier et al., 2018; Metzger et al., 2012; Monroy et al., 2018; Sung et al., 2008). In mammalian myotubes, MAP7 is needed for proper kinesin-1-dependent nuclear distribution (Metzger et al., 2012), but whether MAP7 proteins are needed for other kinesin-1-dependent processes in mammals has not been investigated. It is also unknown whether mammalian MAP7 homologues all behave similarly and whether they have different, overlapping or redundant functions.

**Figure 1.**
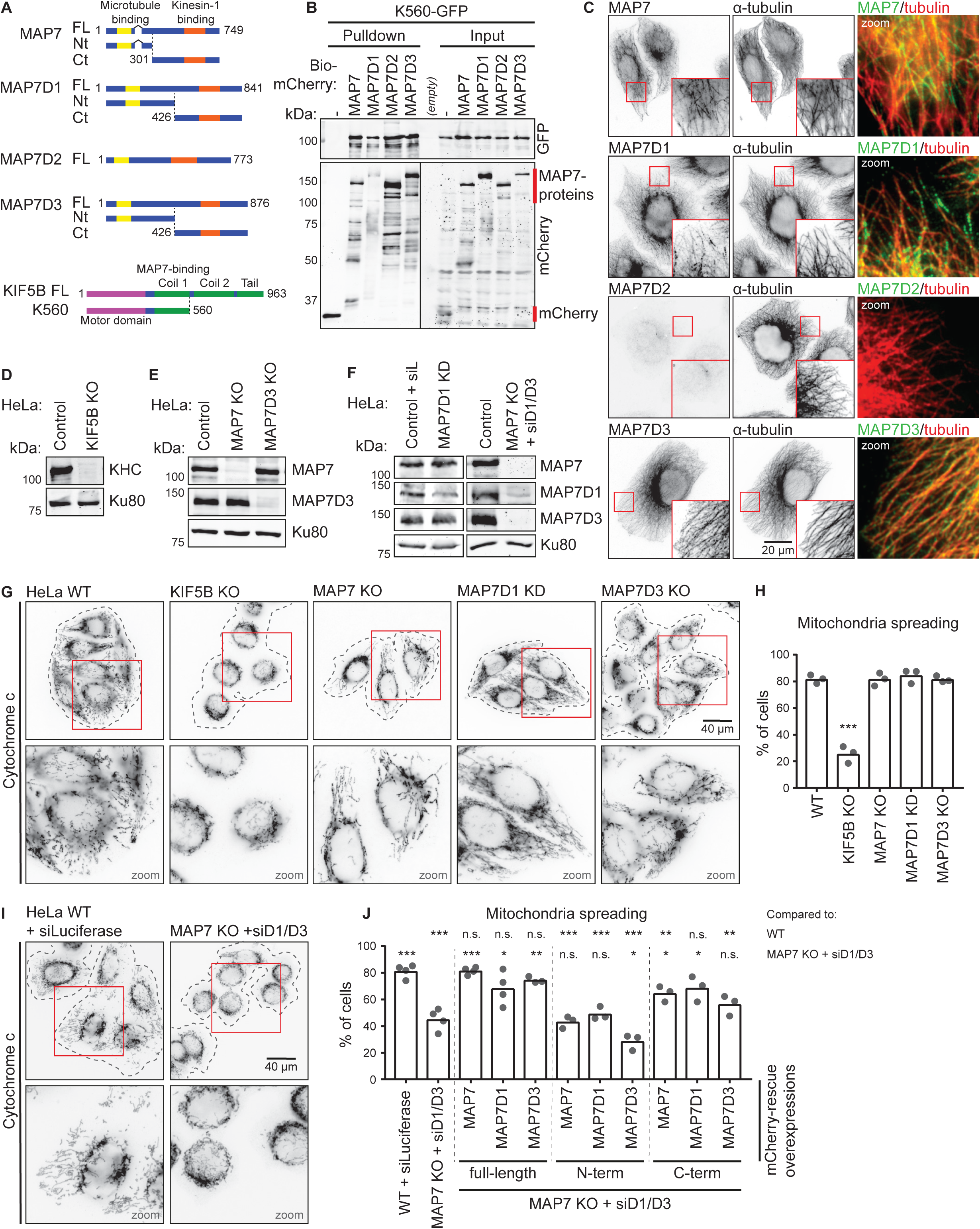
Redundant function of MAP7 family proteins in kinesin-1-dependent mitochondria distribution. (A) Schemes of MAP7 family proteins and KIF5B constructs. (B) Streptavidin pull down assay with extracts of HEK293T cells expressing BirA, K560-GFP (prey) and the indicated Bio-mCherry-labeled proteins (bait) analyzed by Western blotting. Red lines indicate the position of mCherry (negative control) and MAP7 proteins. (C) Immunostaining of HeLa cells for endogenous MAP7 family members and α-tubulin imaged on a wide field microscope. (D-F) Western blot analysis of the indicated HeLa knockout (KO) and knockdown (KD) cells with the indicated antibodies; Ku80 was used as a loading control. (G, I) HeLa cells treated as indicated stained for mitochondria (cytochrome c). Cell outlines are indicated with grey dashed lines; zooms (red squares) are shown below. (H, J) Mitochondria distribution scored per condition using cytochrome c staining, (H) n = 345, 347, 332, 338, and 373 cells from three independent experiments. Wild type (WT) vs KIF5B KO, p = 0.0002, Student’s t test. (J) n = 444 (WT + siLuciferase), n = 589 (MAP7 KO + siMAP7D1/D3) and for rescue conditions on top of MAP7 KO + siMAP7D1/D3: n = 261, 277, 324, 297, 277, 296, 263, 267 and 416 cells all from three or four independent experiments, Student’s t test: *, p<0.05, **, p <0.01 and *** p <0.001.

Interestingly, in vitro experiments in fly ovary extracts have shown that the full-length kinesin-1, but not its minimal dimeric kinesin-1 fragment requires ensconsin for productive interaction with MTs (Sung et al., 2008). In vitro reconstitutions with purified proteins demonstrated that MAP7 recruited kinesin-1 to MTs and somewhat decreased motor velocity but had only a mild effect on kinesin-1 run length (Monroy et al., 2018). Importantly, MAP7 was highly immobile in these assays and was not co-transported with the motor, suggesting that MAP7 affects only the initial recruitment of the kinesin to MTs but has little impact on kinesin-1 movement (Monroy et al., 2018). However, some observations in flies do not agree with this simple model, as it was shown that the C-terminal fragment of ensconsin, which misses the MT binding domain (Sung et al., 2008), significantly rescues kinesin-1-related transport deficiencies in cells lacking ensconsin (Barlan et al., 2013; Metivier et al., 2018). Kinesin-1 is well known to be autoinhibited by its C-terminal cargo-binding domains (Verhey and Hammond, 2009), and it was proposed that ensconsin plays a role in relieving autoinhibition of the kinesin (Barlan et al., 2013). This possibility is in line with the experiments performed in extracts (Sung et al., 2008), but was not yet tested with purified proteins.

Here, we explored the relationship between kinesin-1 activity and mammalian MAP7 proteins. We found that MAP7 family members act redundantly to promote kinesin-1-dependent distribution of mitochondria, as well as MT binding of kinesin-1 KIF5B fragment 1-560 (K560) (Case et al., 1997), which contains the motor domain and the dimerizing stalk with the MAP7-binding site, but not the cargo-binding and autoinhibitory domains. MT recruitment of K560 was rescued not only by full-length MAP7’s, but also by their C-terminal domains, which lacked the major MT binding region. These results were recapitulated using in vitro reconstitution assays with purified proteins, which provided evidence both for MT tethering and allosteric activation of kinesin-1 by MAP7 family members. In agreement with published data, we found that MAP7 was immobile on MTs in vitro (Monroy et al., 2018), whereas MAP7D3 could be observed moving together with K560 motors. In spite of these differences, both MAPs increased not only the recruitment of kinesin-1 to MTs but also its processivity. Such an effect can be explained if the interaction between MAP7’s and kinesin-1 is weak and transient, and this was confirmed by biochemical and imaging experiments. Taken together, our data show that MAP7 proteins redundantly regulate kinesin-1-dependent transport by acting as MT-tethered recruitment factors and activators of this kinesin.

## Results

### MAP7 family members act redundantly in mitochondrial distribution in HeLa cells

To test whether all four MAP7 family members can potentially act as kinesin-1 regulators, we performed a pull down assay and found that all four MAP7 proteins could bind to the kinesin-1 deletion mutant K560 (Fig. 1A, B. Gene expression analysis at the mRNA and protein level indicated that HeLa cells co-express MAP7, MAP7D1 and MAP7D3 (Kikuchi et al., 2018; Syred et al., 2013), and we confirmed these data by antibody staining (Fig. 1C). In contrast, MAP7D2, which is highly expressed in brain tissue (Niida and Yachie, 2011), was not expressed in HeLa cells. To test if all three MAP7’s are required for kinesin-1 function, we initially used the distribution of mitochondria as readout, because it strongly depends on kinesin-1 KIF5B (Tanaka et al., 1998). In the absence of KIF5B, mitochondria were no longer dispersed in the cytoplasm but were clustered around the nucleus (Fig. 1D, G, H). Next, we generated HeLa cells lacking each individual MAP7 family member. In these cells, the expression of the remaining MAP7’s was not altered and no defects in the localization of mitochondria were observed (Fig. 1E-H).

We next attempted to generate a stable triple knockout of MAP7, MAP7D1 and MAP7D3, but such cells were not viable. It is unlikely that this was due to the lack of kinesin-1-mediated transport, as KIF5B knockout cells displayed no apparent growth or proliferation defects, and the two other kinesin-1 isoforms, KIF5A and KIF5C, do not seem to be expressed in HeLa cells (Nagaraj et al., 2011). Although MAP7 was shown to be phosphorylated and thus inactivated during mitosis (McHedlishvili et al., 2018), it is possible that MAP7 proteins still contribute to cell division, as ensconsin is known to participate in spindle formation in flies (Gallaud et al., 2014), and MAP7D3 was reported to modulate the recruitment of kinesin-13 to the mitotic spindle (Kwon et al., 2016). In order to remove all three MAP7 homologues simultaneously, we performed siRNA-mediated knockdown of MAP7D1 and MAP7D3 in the stable MAP7 knockout line, and this approach resulted in an efficient loss of all three MAP7 family members (Fig. 1F). Depletion of all three MAP7 homologs mimicked the effect of KIF5B knockout, leading to a strong perinuclear clustering of mitochondria (Fig. 1I, J. To exclude that this phenotype was caused by defects in MT organization, we assessed it by antibody staining and found that the overall MT arrangement and density were similar (Fig. S1A, B). Furthermore, live imaging of EB3-GFP showed no differences in MT plus-end growth, and polymerizing MT ends still reached the cell periphery (Fig. S1C-E). We conclude that MAP7 family members act redundantly in mitochondria localization and that this effect is unlikely to be due to alterations in MT network architecture.

Mitochondrial positioning in these cells was rescued by re-expressing individual full length MAP7 proteins, and it was also partially rescued by expressing the C-termini of MAP7 and MAP7D1 (Fig. 1J). Rescue with the MAP7D3 C-terminus was less efficient, because the construct was mostly accumulated in the nucleus, and, as its concentration in the cytoplasm was low, only highly expressing cells showed rescue (Fig. S1F).

Rescue of kinesin-1 function by MAP7 C-termini could be potentially explained by their residual affinity for MTs. For example, recent work has demonstrated that MAP7 contains an additional MT binding domain (termed “P-region”) within the intrinsically disordered linker part of the protein (Tymanskyj et al., 2018) (Fig. 2A). To address this possibility, we systematically examined the ability of different parts of MAP7 linker to interact with MTs (Fig. 2A). Using a MT pelleting assay, we found that the C-terminal MAP7 fragment used in cellular experiments (Fig. 1A, J) did not co-sediment with MTs (Fig. 2B and S1G). We were also unable to detect the binding of the mCherry-tagged version of this fragment to MTs in vitro (Fig. S1H). Next, we tested MT binding of different MAP7 fragments by overexpression of their GFP-tagged fusions. Although we could re-confirm MT affinity of the P-region, its smaller fragments as well as other MAP7 deletion mutants lacking the N-terminal MT binding domain, including the MAP7-Ct and Ct(mini), showed no MT enrichment in HeLa cells (Fig. 2A, C and D). However, we could detect weak MT binding of the proline-rich part of the MAP7 linker as well as MAP7-Ct(mini) in COS7 cells, likely due to their flat morphology and low MT density (Fig. 2D and S2A). These data suggest that the C-terminal part of the MAP7 linker might have some weak MT affinity, which could contribute to but is unlikely to fully explain the ability of MAP7 C-terminus to rescue KIF5B-dependent mitochondria localization.

**Figure 2.**
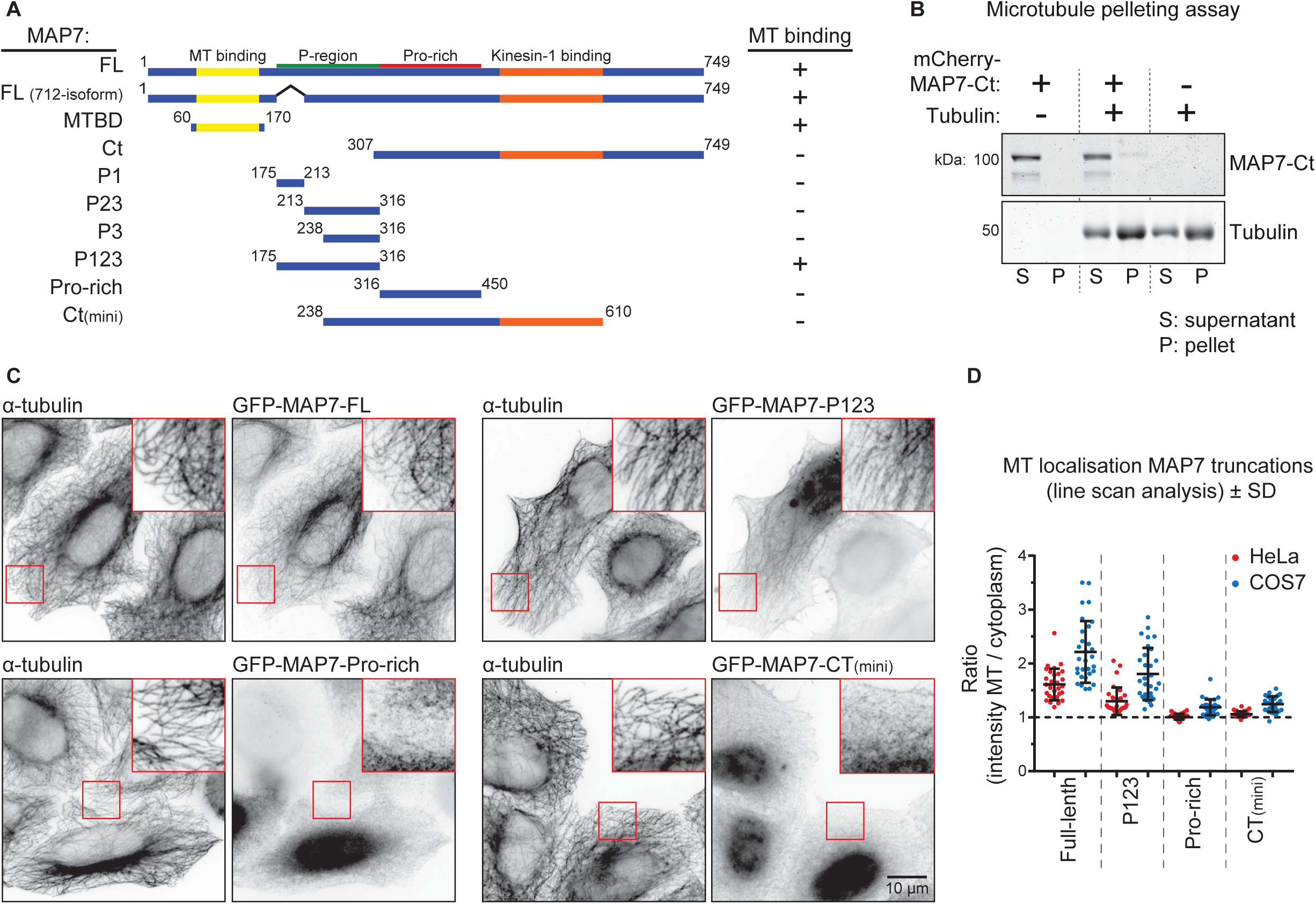
Characterization of MT binding domains of MAP7. (A) Scheme of MAP7 truncations. MT binding was assayed by overexpression of GFP-tagged constructs and co-stain for α-tubulin; examples are shown in panel C. (B) MT pelleting assay with mCherry-MAP7-Ct analyzed by SDS-PAGE. Uncropped gel images are shown in Fig. S1G. (C,D) Indicated GFP-tagged MAP7 constructs were overexpressed in MAP7 KO HeLa cells co-stained for α-tubulin (C), to quantify their MT enrichment by line scan analysis (D). n = 10 cells representing 30 MTs (3 per cell) per condition.

### K560 binding to MTs in cells depends on MAP7 proteins

To show that the loss of MAP7 proteins has a direct effect on kinesin-1 activity, we next examined the distribution of the dimeric K560 truncation mutant that can move along MTs but does not bind to cargo. In control HeLa cells, this construct was distributed along MTs, and in most cells, it showed enhanced accumulation on MTs in cell corners, where MT plus ends are concentrated (Fig. 3A). Depletion of individual MAP7 family members did not alter this distribution except for the knockout of MAP7D3, in which less K560 accumulated on corner MTs (Fig. 3A, B. In contrast, in cells lacking all three MAP7 proteins, K560 showed a diffuse localization (Fig. 3C, E. Expression of MAP7, MAP7D1 or MAP7D2 in such cells rescued the recruitment of the kinesin to MTs, whereas expression of MAP7D3 led to strong co-accumulation of both constructs on MTs in the corners of almost all transfected cells (Fig. 3D-F). Expression of the N-terminal, MT binding fragments of MAP7 and its homologs could not restore the distribution of K560, whereas significant rescue of MT binding by the kinesin was observed with the C-termini of all MAP7 proteins (Fig. 2E and S2B). We conclude that K560 displays low binding to cellular MTs in the absence of MAP7, and that this binding can be increased by kinesin-1-interacting C-terminal MAP7 fragments, which are diffusely localized in HeLa cells. The C-terminus of MAP7D3, which was mostly nuclear on its own (Fig. S1F), was retained in the cytoplasm when expressed together with K560, and shifted the localization of this kinesin fragment to MTs in cell corners, similar to the full-length MAP7D3 (Fig. 3D, E and S2B).

**Figure 3.**
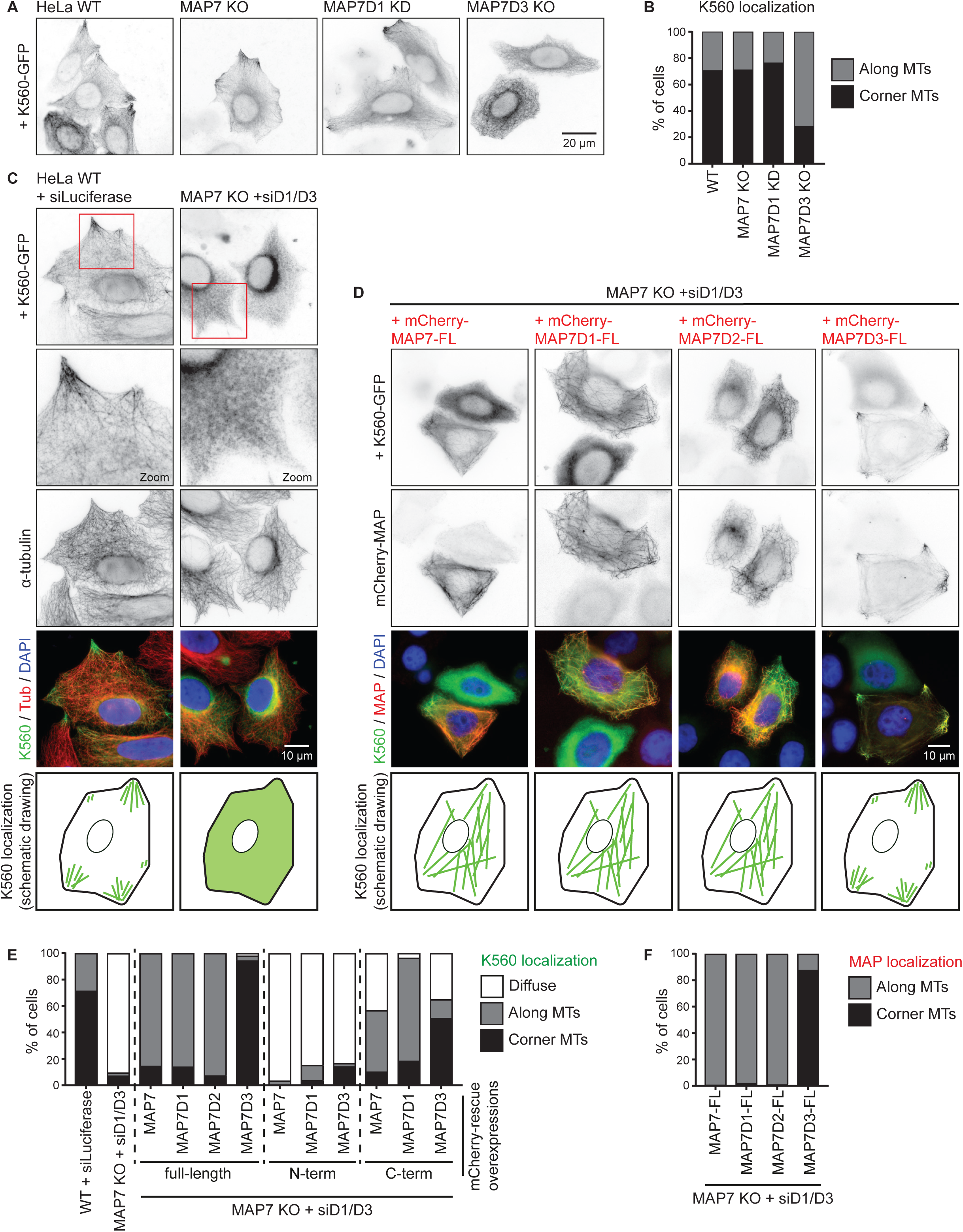
Kinesin-1 recruitment to MTs depends on MAP7 family proteins. (A,B) Wide field images of K560-GFP overexpressed in the indicated HeLa control, KO or KD conditions (A), and quantification of K560-GFP localization (B). n = 234, 252, 174, and 254 cells from three independent experiments. (C,D) Wide field images of K560-GFP overexpressed either alone or together with mCherry-tagged MAP7 constructs in control or MAP7 KO + siMAP7D1/D3 HeLa cells, as indicated. In (C), cells were co-stained for α-tubulin. A schematic drawing of K560-GFP localization is shown at the bottom. (E,F) Quantification of K560-GFP (E) and MAP7 construct (F) localization per condition, as indicated, categorized as: diffuse, along MTs or at corner MTs. n = 459 (WT + siLuciferase), n = 485 (MAP7 KO + siMAP7D1/D3) and for rescue conditions: n = 113, 237, 90, 167, 210, 186, 133, 176, 197 and 193 cells from two to four independent experiments.

### MAP7D3 but not MAP7 can be redistributed by kinesin-1

To understand why MAP7D3 but not the other MAP7 homologues promotes MT plus-end shifted distribution of K560, we next examined the distribution of endogenous MAP7 and MAP7D3 and found that only MAP7D3 could be efficiently relocalized by K560 to cell corners (Fig. 4A, B).

**Figure 4.**
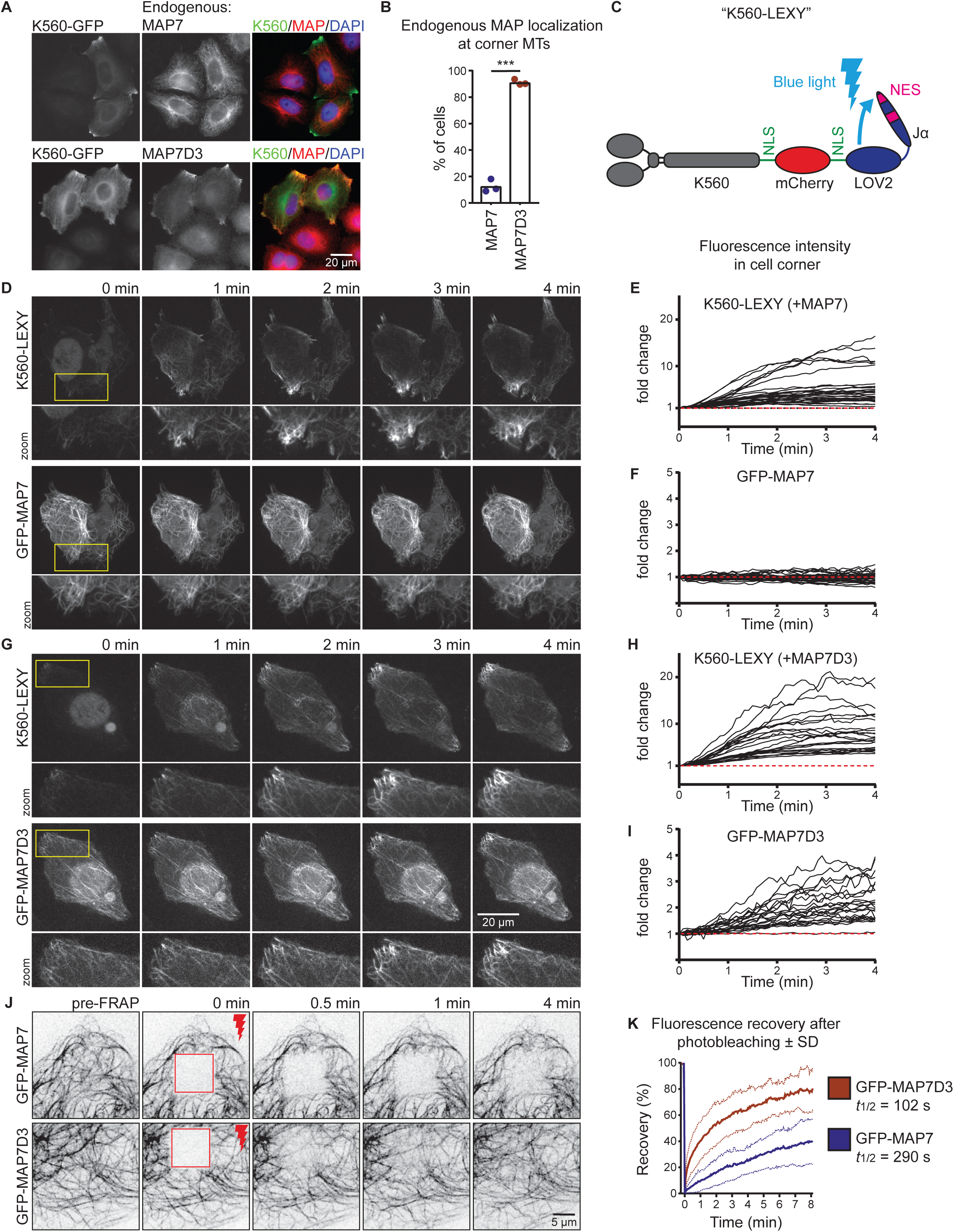
Kinesin-1 can redistribute MAP7D3 in cells. (A-B) Wide field images of K560-GFP overexpressed in HeLa cells stained for endogenous MAP7 or MAP7D3 (A) used to quantify endogenous MAP localization (B). n = 232 and 303 cells from three independent experiments, Student’s t test: p < 0.001. (C) A scheme of K560-LEXY construct containing two NLS sequences and an mCherry tag. Blue light induces a conformation change, causing detachment of the Jα-peptide containing a nuclear export signal from the LOV2 domain. (D, G) Single frames of KIF5B KO cells co-transfected with K560-LEXY and GFP-MAP7 (D) or GFP-MAP7D3 (G) sequentially illuminated with green and blue light (in that specific order). Zooms are indicated in yellow. (E, F, H, I) Measurements of fluorescence intensity changes over time in K560-LEXY-positive cell corners. Black lines represent single measurements of K560-LEXY (E,H), GFP-MAP7 (F) and GFP-MAP7D3 (I). n = 31 measurements from 17 cells (E, F) and n = 22 measurements from 14 cells (H, I), from two independent experiments. (J) Single frames of FRAP experiments on COS7 cells overexpressing GFP-MAP7 or -MAP7D3. Stills show a baseline (pre-FRAP), the first frame after photobleaching (0 min) and the indicated time points after FRAP of a 10×10 μm square region (shown in red). (K) Quantification of fluorescence recovery of J. The graph shows mean curves (bold lines) ±SD (light dotted lines) over time. n = 18 cells from three independent experiments for each condition.

To prove that kinesin-1 can indeed rapidly relocalize MAP7D3, we have set up an optogenetics-based assay, in which K560 could be sequestered in the nucleus and then acutely released from it using a light-inducible nuclear export system (Niopek et al., 2016). A K560-mCherry, containing NLS sequences, was C-terminally tagged with an engineered domain of *Avena sativa* phototropin-1, AsLOV2, in which the Jα helix was modified to contain a nuclear export signal (Fig. 4C). Within 1 to 2 minutes after activation with blue light, K560 was efficiently exported from the nucleus (Fig. 4D, E, G, H). MAP7D3, but not MAP7 co-accumulated on MTs in cell corners within a 4-minute time frame (Fig. 4D-I, Video 1 and 2). We conclude that K560 can indeed acutely relocalize its own positive regulator MAP7D3, but not MAP7, when kinesin expression is sufficiently high.

To explain why the distribution of MAP7D3 but not that of MAP7 was sensitive to the presence of kinesin-1, we hypothesized that MAP7D3 might be more mobile on MTs. To test this idea, we performed Fluorescence Recovery after Photobleaching (FRAP) experiments with GFP-tagged MAP7 and MAP7D3 and found that the latter indeed exchanged much more rapidly on MTs (Fig. 4J, K. The different turnover rates of the two MAP7 family proteins on MTs, possibly combined with the different affinities to kinesin-1, seem to contribute to their differential relocalization by overexpressed kinesin-1.

### MAP7 and MAP7D3 control kinesin-1 recruitment to MTs and motor processivity

To get further insight into the similarities and differences in the regulation of kinesin-1 by MAP7 proteins, we set up in vitro reconstitution assays. In contrast to previously published experiments, which employed taxol-stabilized MTs in the absence of free tubulin, we used dynamic MTs that were grown from GMPCPP-stabilized seeds (Bieling et al., 2007). Kinesins, MAPs and MTs were observed by Total Internal Reflection Fluorescence Microscopy (TIRFM), as described previously (van Riel et al., 2017). To study kinesin-1 motility, we purified full-length KIF5B-GFP and K560-GFP from HEK293T cells (Fig. S3A). Analyses by mass spectrometry and Western blotting showed that although some co-purification of MAP7, MAP7D1 and MAP7D3 with this kinesin was observed when the protein was washed with a low ionic strength buffer, this contamination was almost entirely removed when the ionic strength of the washing buffer was increased (Fig. S3B-E). We used such “high-salt washed” kinesin preparations for all our experiments. In cells, kinesin-1 can exist in a heterotetrametric form with two light and two heavy chains. Although some light chains could be detected by Western blotting (Fig. S3B), we assume that in our assays most full-length kinesins were dimers of heavy chains as no light chains were visible on a Coomassie-stained gel (Fig. S3A, red arrow).

The MAP7 binding site on kinesin-1 is well defined (Monroy et al., 2018) and pull down assays confirmed that there are no additional binding sites for MAP7 in the C-terminal coil or the tail of kinesin-1 (Fig. 5A, B. Kymograph analysis of full length kinesin-1 motors showed two populations of single molecule behavior: short binding events which did not result in processive movement (classified as events that last ≤1 second) or events of landing followed by processive movement. Addition of purified Alexa647-labeled SNAPtag-MAP7 or -MAP7D3 (Fig. S3A) in these assays led to a dramatic increase (at least 60 fold) of both type of events (Fig. 5C, D. Furthermore, processive kinesin runs were on average ~1.6 fold longer (Fig. 5C, E, while the velocities were, especially upon addition of MAP7D3, reduced (Fig. 5C, F and S3G).

**Figure 5.**
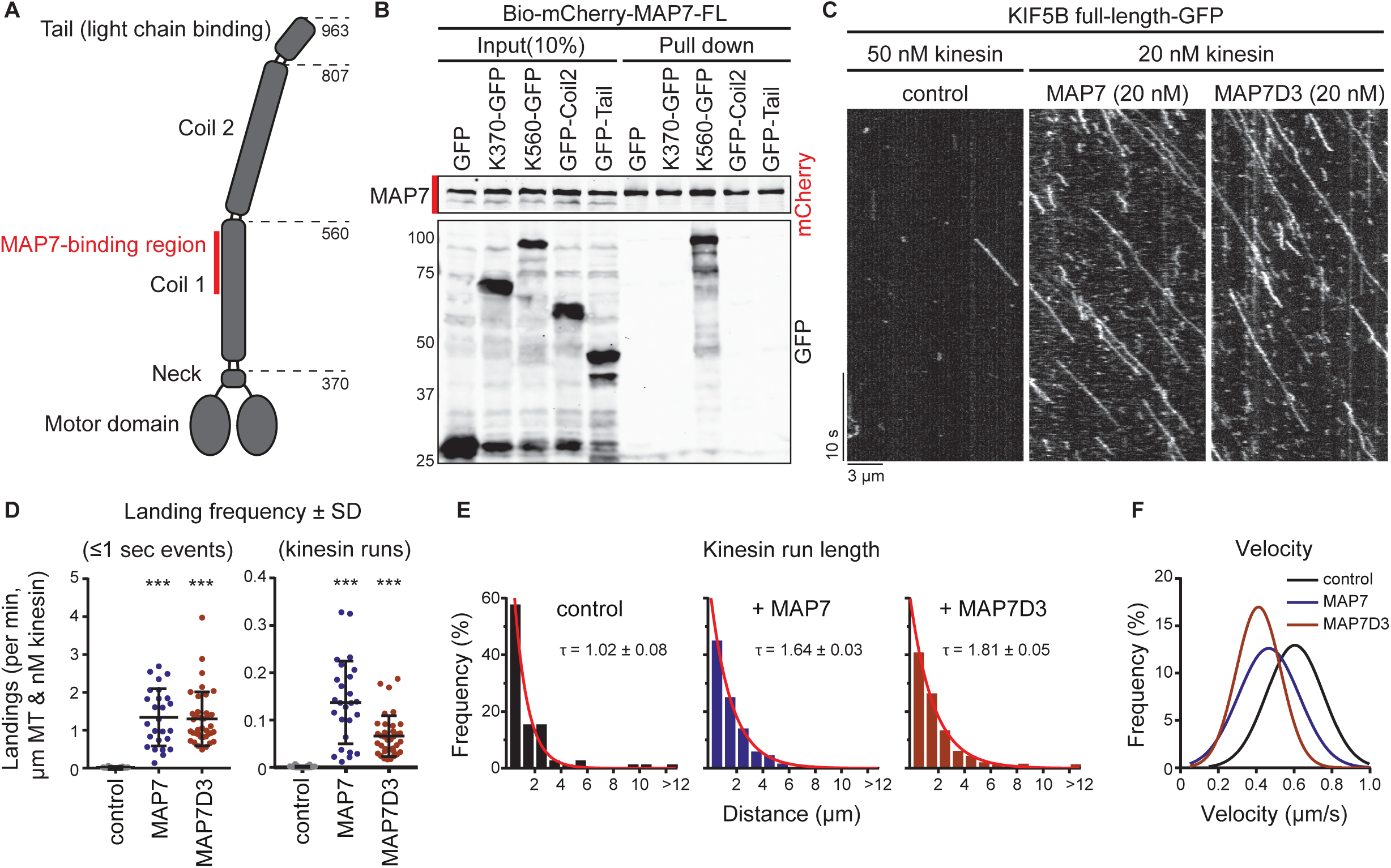
Kinesin-1 is regulated by MAP7 proteins in vitro. (A) Overview of full-length kinesin-1 (KIF5B) domains. (B) Streptavidin pull down assay with extracts of HEK293T cells over-expressing BirA, the indicated KIF5B-GFP truncations (prey) and Bio-mCherry-MAP7 (bait) analyzed by Western blotting. (C) Kymographs of GFP-tagged full-length KIF5B (kinesin-1) on dynamic MTs in control conditions or in the presence of MAP7 or MAP7D3. (D) Quantification of kinesin-1 landing frequencies per MT and corrected for MT length, time of acquisition and kinesin concentration. n = 99, 26 and 38 MTs from 2 or 3 independent experiments. (E) Histograms of run lengths fitted to a single exponential decay (red) with indicated rate constants (tau) as a measure of mean run length. n = 71, 542 and 568 kinesin runs from 2 or 3 independent experiments. (F) Gaussian fits of kinesin velocities. Histograms are shown in Fig. S3G.

To further examine the effects of MAP7 and MAP7D3 on kinesin-1, we turned to the K560 fragment, which contains the MAP7-binding site but lacks the cargo-binding and autoinhibitory tail. When K560 was added to MAP7 or MAP7D3-decorated MTs, we observed a strong (up to 23.6 fold) increase in the motor landing frequency compared to K560 alone (Fig. 6A and B), in agreement with published data on MAP7 (Monroy et al., 2018). The landing frequency of K560 increased with higher MAP concentrations and correlated with increasing MT labeling intensity by the particular MAP (Fig. 6A-C and S4A). Furthermore, we found that MAP7D3 but not MAP7 caused a very significant decrease in kinesin velocity (Fig. 6D and S4B). Finally, we found that both MAP7 and MAP7D3 could induce a 2-fold increase in kinesin processivity (Fig. 6E-G), with some kinesin runs exceeding 10 µm in length. We note that for this quantification, we only took into account the runs, in which we observed both kinesin association and dissociation from the MT. Inclusion of all detected runs suggested that in the presence of MAP7 or MAP7D3, even longer runs could occur (not shown).

**Figure 6.**
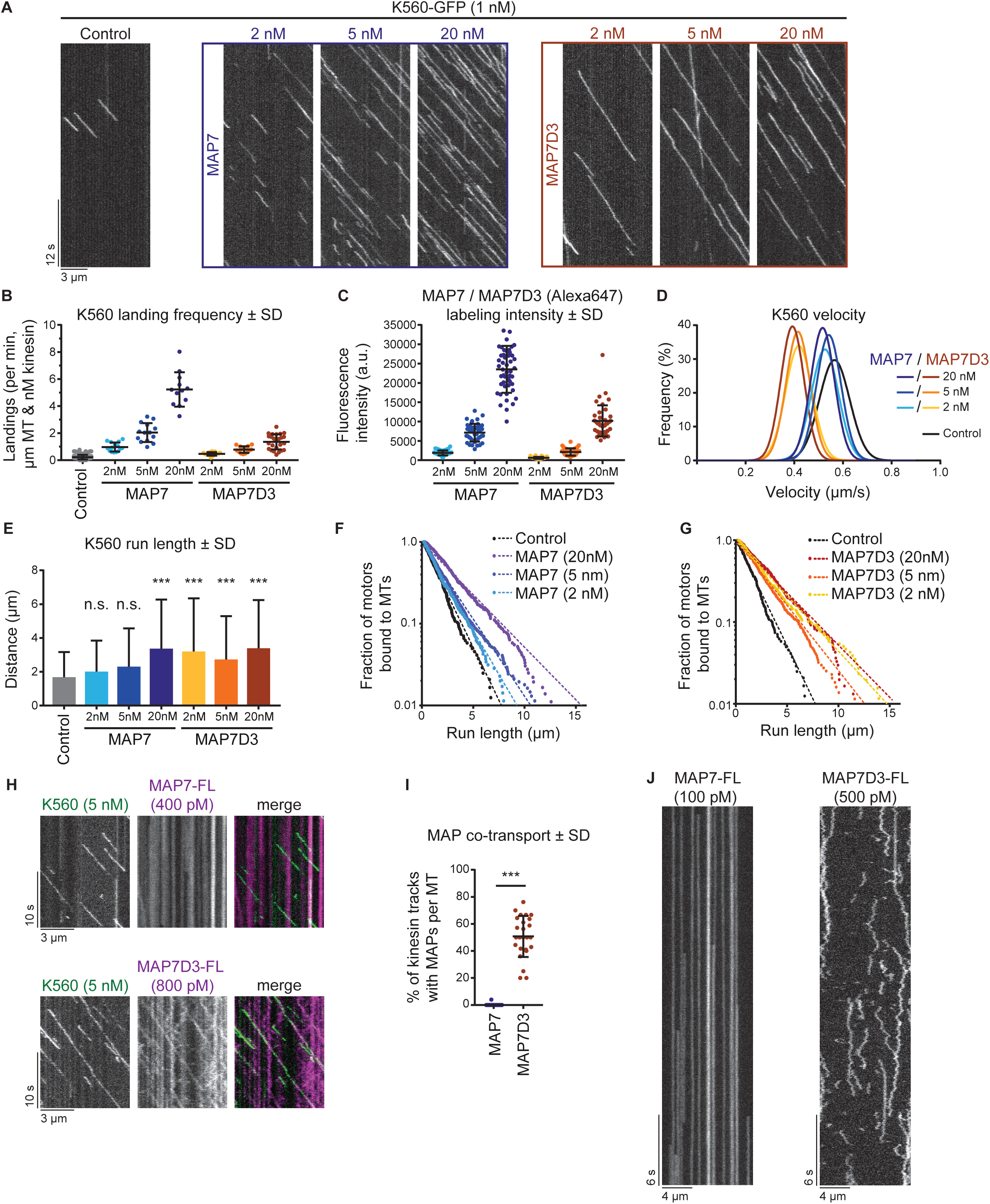
Density and mobility of MAP7’s determine kinesin-1 landing and processivity. (A) Kymographs of K560-GFP on dynamic MTs in control conditions or in the presence of increasing concentrations of MAP7 or MAP7D3. (B) Quantification of kinesin landing frequencies per MT and corrected for MT length, time of acquisition and kinesin concentration. n = 167, 15, 14, 12, 13, 15 and 25 MTs from two independent experiments. (C) Quantification of SNAP(Alexa647)-MAP7 and -MAP7D3 intensities on dynamic MTs using images acquired under identical conditions on a TIRF microscope. n = 39 to 49 MTs from two independent experiments. Representative images are shown in Fig. S4A. (D) Gaussian fits of kinesin velocities. Histograms are shown in Fig. S4B. (E) Quantification of K560-GFP run length. n = 241, 351, 614, 361, 257, 436 and 303 kinesin runs from two independent experiments. Mann-Whitney U test: ***, p <0.001. (F, G) Cumulative distributions of K560-GFP run lengths measured in the presence of increasing concentrations of MAP7 (F) or MAP7D3 (G). Straight dashed lines correspond to single exponential fits, n numbers correspond to panel E. (H) Kymographs of dual-color in vitro reconstitution experiments with K560-GFP and SNAP(Alexa647)-tagged MAP7 or MAP7D3. (I) Quantification of kinesin tracks positive for MAP7 or MAP7D3 co-transport, n = 417 from 19 MTs (MAP7) and n = 344 from 25 MTs (MAP7D3) from two independent experiments, Mann-Whitney U test: ***, P <0.001. (J) Kymograph of single SNAP(Alexa647)-MAP7 or -MAP7D3 molecules on dynamic MTs in vitro. Movies were acquired at 25 frames/sec on a TIRF microscope.

The increase of run lengths in the presence of MAP7 and MAP7D3 could be explained by kinesin multimerization or by a model where MAP7 acts as an additional MT attachment point. In these cases the distribution of run lengths is expected to be described by the sum of two or three exponential decays (Klumpp and Lipowsky, 2005). However, the corresponding best fit of distributions in Fig. 6F, G converged to a single exponential decay, suggesting that MAP7 directly affects kinesin’s binding/unbinding rate constants, instead of introducing an additional intermediate binding state. Moreover, single molecule analysis of K560 moving on MTs showed that kinesin-1 intensity profiles matched that of a single dimer in assays both with and without MAP7D3 (Fig. S4C). In addition, we performed mixed kinesin assays where GFP- and SNAP(Alexa647)-tagged kinesins were used in a 1:1 ratio. If kinesin-1 would multimerize in the presence of MAP7 proteins, then one would expect to see a significant fraction of two-colored kinesin tracks per kymograph; however, such events were not observed (Fig. S4D), confirming our observation of K560 behaving as a single dimer on MAP7-decorated MTs. Taken together, these data suggest that the presence of MAP7 alters the state of single kinesin dimers.

Interestingly, MAP7 could promote kinesin processivity at high concentrations, when MTs were fully decorated, whereas MAP7D3 reduced kinesin detachment from MTs even at low concentrations (Fig. 6C, E-G). These data correlated with the observation that MAP7D3 but not MAP7 could move together with K560 in vitro (Fig. 6H, I. To find explanation for this difference in co-transport we examined single molecule dynamics of MAP7 and MAP7D3 on in vitro polymerized MTs and found that MAP7 showed very long static binding events, many of which exceeded our observation time (5 min) (Fig. 6J), in agreement with recently published data (Monroy et al., 2018). In contrast, MAP7D3 displayed a diffusive behavior, with many short binding events (Fig. 6J). These data are in agreement with the FRAP data, showing that in cells, MAP7D3 is more mobile than MAP7 (Fig. 4J, K. The density of MT labeling was higher with MAP7 than with MAP7D3 at the same protein concentration, indicating that the latter has a lower affinity for MTs (Fig. 6C, J and S4A). We conclude that MAP7 proteins can affect not only kinesin landing on MTs, as suggested previously (Monroy et al., 2018; Sung et al., 2008), but its processivity. Co-transport of the MAP with the kinesin could facilitate processive motion, but was not essential, as also a statically bound MAP7 could exert this effect if its density on MTs was high enough.

### MAP7D3 C-terminus promotes MT recruitment and processivity of kinesin-1 in spite of having only a low MT affinity

The ability of K560 to transport MAP7D3 but not MAP7 might be caused not only by the different MT binding behavior of the two MAPs, but also by their different affinities for the kinesin. To test this possibility, we purified minimal binding constructs for MAP7, MAP7D3 and KIF5B from *E. coli* (Fig. 7A and Fig. S3F) and probed the oligomerization state of the individual proteins in solution by size exclusion chromatography followed by multi-angle light scattering (SEC-MALS). As expected, both the MAP7 and MAP7D3 fragments were monomers with the measured molecular weight (MW) values of 18.4 kDa (calculated MW of 18.8 kDa) and 22.2 kDa (calculated MW of 21.5 kDa), respectively, whereas the KIF5B fragment was a dimer (measured and calculated MW of 25.9 kDa) (Fig. 7B). Using isothermal titration calorimetry (ITC) experiments, we found an equilibrium dissociation constant K_D_ of 3.8 ± 0.2 µM and a stoichiometry number, N, of 0.96 (monomer equivalents) for the interaction between the KIF5B and MAP7D3 fragments (Fig. 7C and S4E, F). In contrast, the MAP7 fragment had an affinity for the KIF5B fragment that was too weak to be properly determined; however, the obtained isotherm suggested a K_D_ in the higher micromolar range (Fig. 7D and S4E, G).

**Figure 7.**
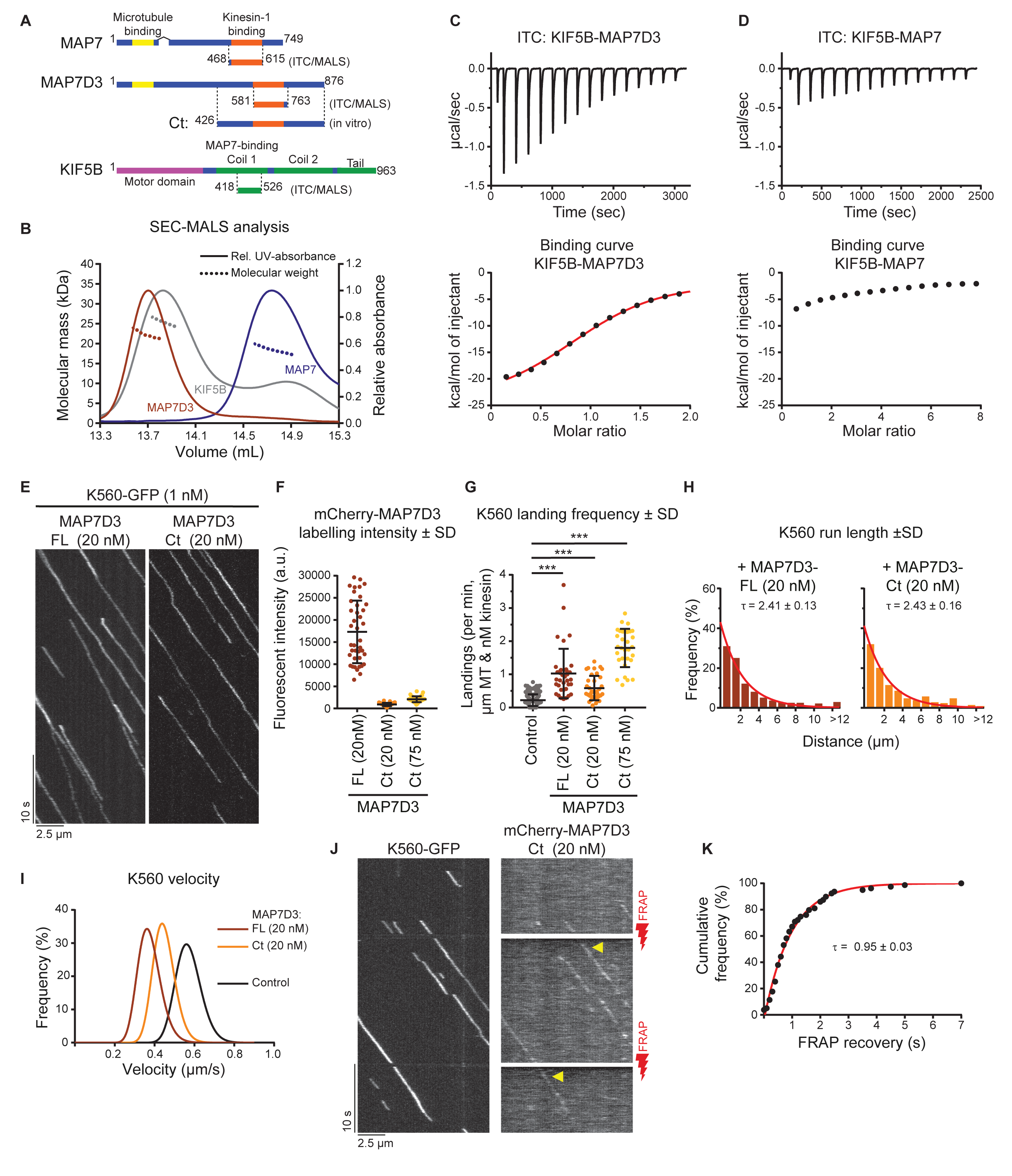
MAP7D3 binds to KIF5B more tightly than MAP7 and promotes its activity in vitro. (A) Schemes of MAP7 family proteins and KIF5B constructs used in this figure. (B) SEC-MALS was used to determine the oligomerization state of indicated proteins in solution. Lines depict the relative UV-absorbance measured at 280 nm plotted against the elution volume. The molecular weight determinations by multi-angle light scattering are depicted by dashed lines. (C, D) ITC of MAP7D3 (C) or MAP7 (D) against KIF5B. Top panels: Enthalpograms of the respective titrations. Lower panel: integrated heat change (black dots) and the associated fitted curve (red line in C). Controls are shown in Fig. 4A-C. (E) Kymographs of K560-GFP on dynamic MTs in the presence of full-length (FL) MAP7D3 or MAP7D3-Ct (both purified from *E.coli*). (F) Quantifications of MAP7D3-FL or -Ct intensities on dynamic MTs using images acquired under identical conditions on a TIRF microscope. n = 40, n= 38 and n= 42 MTs from two independent experiments. Representative images are shown in Fig. S5A. (G) Quantification of landing frequency per MT and corrected for MT length, time of acquisition and kinesin concentration. n = 167, 36, 36 and 31 MTs from two independent experiments. (H) Histograms showing kinesin run lengths fitted to a single exponential decay (red) with the indicated rate constants (tau) as a measure of mean run length, n = 241, 271 and 209 kinesin runs from two independent experiments; the associated bar plots are shown in Fig. S5C. (I) Gaussian fits of kinesin velocities. Histograms are shown in Fig. S5B. (J) Kymographs of single molecule FRAP experiments on K560-GFP motors and MAP7D3-Ct. Photobleaching with a 561 nm red laser was performed at time points indicated with red lightning bolts. Fluorescent recovery is indicated with a yellow arrow. (K) Cumulative frequency distribution plot of mCherry-MAP7D3-Ct recovery after photobleaching (black dots) fitted to an exponential decay (red line) with the indicated decay constant (tau), n = 79 from three independent experiments.

Since MAP7D3 seems to be the most potent kinesin-1 interactor, we set out to compare the effect of full-length MAP7D3 and its C-terminus (Ct, also used for cellular experiments), both purified from *E. coli* (Fig. 7A and Fig. S3F), on K560 motility in vitro (Fig.7E). In agreement with a previous publication (Yadav et al., 2014), MAP7D3-Ct displayed weak MT binding: MT labeling intensity with 20 nM MAP7D3-Ct was 19.2 fold lower than with 20 nM full length MAP7D3 (Fig. 7F and S5A). In spite of this lower MT affinity, MAP7D3-Ct could efficiently increase K560 landing rate, promote its processivity and decrease motor velocity compared to control (Fig. 7E-I and S5A-C). The effect of MAP7D3-Ct on the landing rate was particularly obvious at 75 nM concentration, as MT labeling at this concentration was still 8.3 fold lower than with 20 nM full length MAP7D3, whereas the K560 landing frequency was 8.1 fold higher compared to control (kinesin only) (Fig. 7F,G). These data argue against the simple model that MAP7D3 acts as MT-recruiting factor for kinesin-1, but cannot exclude that the weak binding of MAP7D3 C-terminus to MTs augments K560-MT interaction. These data also show that the addition of a weak MAP module to the kinesin coil is sufficient to make kinesin-1 processive.

Simultaneous imaging of K560-GFP and mCherry-MAP7D3-Ct showed that this truncated MAP colocalized with moving kinesin (Fig. 7J). Importantly, FRAP analysis in the mCherry channel, leaving the GFP fluorescence unaffected, showed that MAP7D3-Ct exchanged rapidly on moving K560 motors (Fig. 7J, K. Together with the micromolar-range K_D_ of the kinesin-MAP binding (Fig. 7C, D, these data support the idea that fast binding-unbinding kinetics enables static MAP7 proteins to promote processive movement of kinesin-1.

### MAP7 C-terminus promotes kinesin-1 landing on MTs independently of MT tethering

To investigate whether MAP7 family proteins can exert an effect on kinesin-1 that is independent of MT tethering, we used MAP7-Ct(mini) (Fig. 8A), which was completely diffuse in HeLa cells, showed only very weak MT binding in COS7 cells (Fig. 2A,C,D and S2A) and could be prepared from *E. coli* at high concentration in an untagged form (Fig. S3F). When added at micromolar concentrations to the assay with K560, this protein fragment caused a significant (3.4 fold) increase in the motor landing frequency (Fig. 8B, C, whereas the velocity of the kinesin was only mildly affected (Fig. S5E). Strikingly, the increase in motor processivity observed with the MAP7D3 C-terminus was not detected with MAP7-Ct(mini) (Fig. S5F, G).

**Figure 8.**
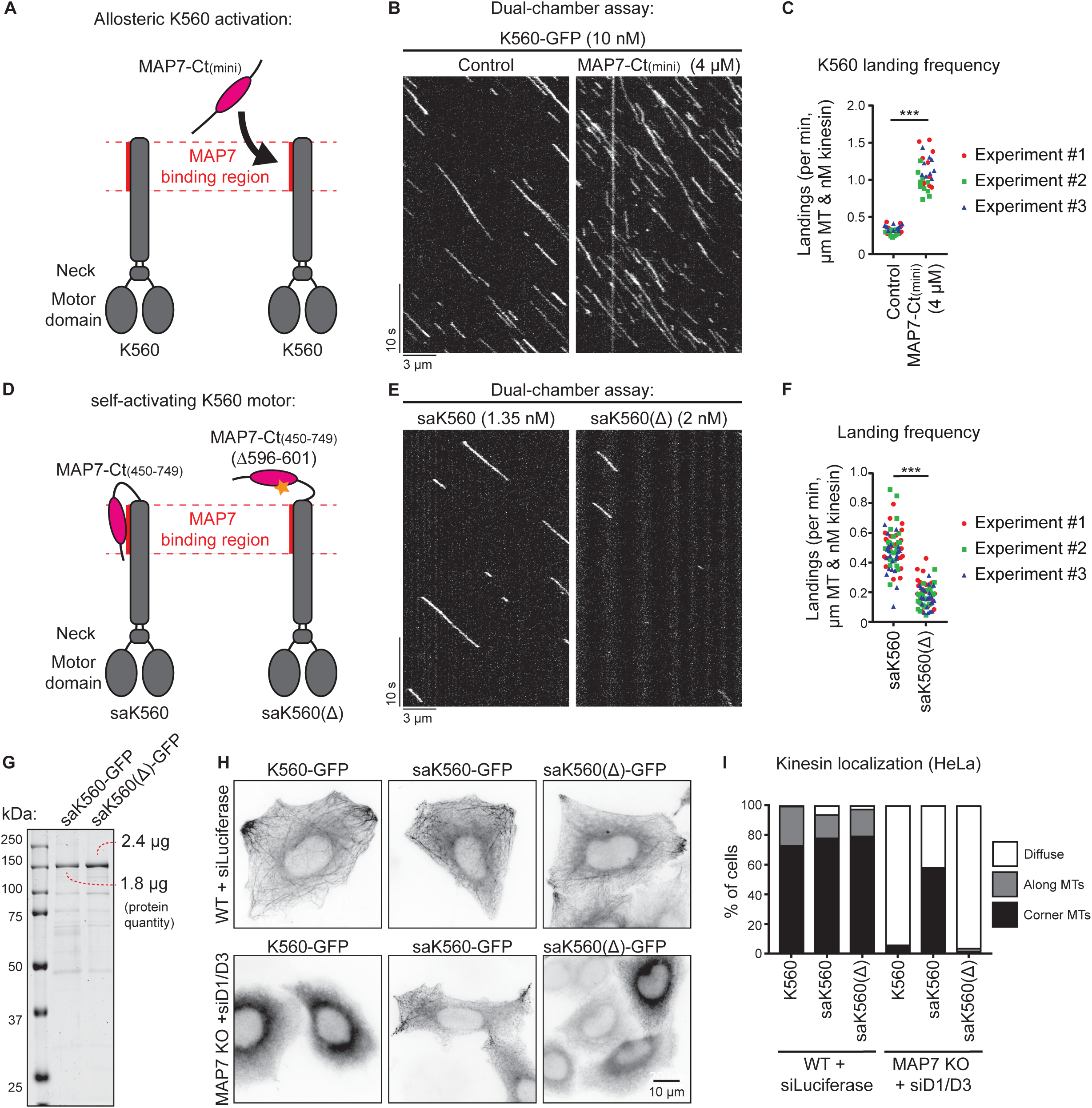
MAP7 C-terminus can activate kinesin-1. (A,D) Overview of kinesin-1 and MAP7 constructs used for experiments. (B,E) Kymographs of a dual-chamber in vitro experiment, where equal concentrations of K560-GFP motors where added to chambers with or without MAP7-Ct(mini) (B), or with the indicated concentrations of saK560/ saK560(∆) (E) on dynamic MTs. (C,F) Landing frequencies quantified per MT and corrected for MT length, time of acquisition and kinesin concentration. Each independent dual-chamber experiment is color-coded, n = 32 and 29 MTs (C) and n = 73 and 69 MTs (F), all from three independent experiments, Mann-Whitney U test: ***, p <0.001. (G) Analysis of purified saK560 proteins by SDS-PAGE. Protein concentrations were determined from a single gel using BSA standard. (H) Wide field images of overexpressed GFP-tagged kinesin constructs in control or MAP7 KO + siMAP7D1/D3 HeLa cells. (I) Quantification of kinesin localization categorized as: diffuse, along MTs or at corner MTs. n = 203, 172, 198, 241, 186 and 234 cells from three independent experiments.

To further substantiate the observation that the MAP7 C-terminus can activate kinesin-1 independently of any residual MT affinity, we developed a self-activating K560 motor, saK560, where the kinesin-binding domain of MAP7 lacking both the P-region and the Pro-rich region was fused to the C-terminus of K560. As a negative control, we generated a saK560(Δ) mutant lacking six amino acids of MAP7 essential for the kinesin binding (Fig. 8D) (Monroy et al., 2018). Comparison of these two kinesins showed that the saK560 had a higher landing frequency (Fig. 8E-G), while other parameters such as velocity and run length were not much affected (Fig. S5H-J). In further support of this observation, we examined the behavior of K560 together with the self-activating fusion proteins in HeLa cells and saw that they all behaved very similarly in wild type cells (Fig. 8H, top). However, upon depletion of all MAP7 family proteins, saK560(Δ) motor was diffuse, just like K560; yet, the saK560 kinesin showed enhanced localization on MTs in cell corners (Fig. 8H, I, demonstrating that this motor still displays activity despite of the absence of MAP7 family protein. Altogether, MT landing of kinesin-1 can be increased by MAP7 family proteins independently of their MT interaction, whereas the regulation of kinesin processivity by these MAPs depends on their association with MTs.

### The stalk of K560 inhibits MT interaction

Our finding that MAP7 C-terminus improves the MT landing frequency of K560 could potentially be explained if the MAP7-interacting stalk of kinesin-1 partly interferes with MT binding. If this were true, a kinesin-1 truncation lacking this stalk should bind to MTs more efficiently. To test this idea, we generated a shorter KIF5B truncation mutant, KIF5B 1-370 (K370), which lacks the MAP7-binding coil region but still dimerizes via its neck linker (Kozielski et al., 1997) (Fig. 9A,B and Fig. S5D). In in vitro assays, K370 appeared to be a faster kinesin with slightly shorter runs compared to K560 (Fig. 9C and S5K-M). Importantly, we observed a 7.8 fold difference in motor landing frequency (Fig. 9D), indicating that the presence of the stalk region in K560 has a negative effect on its interaction with MTs.

**Figure 9.**
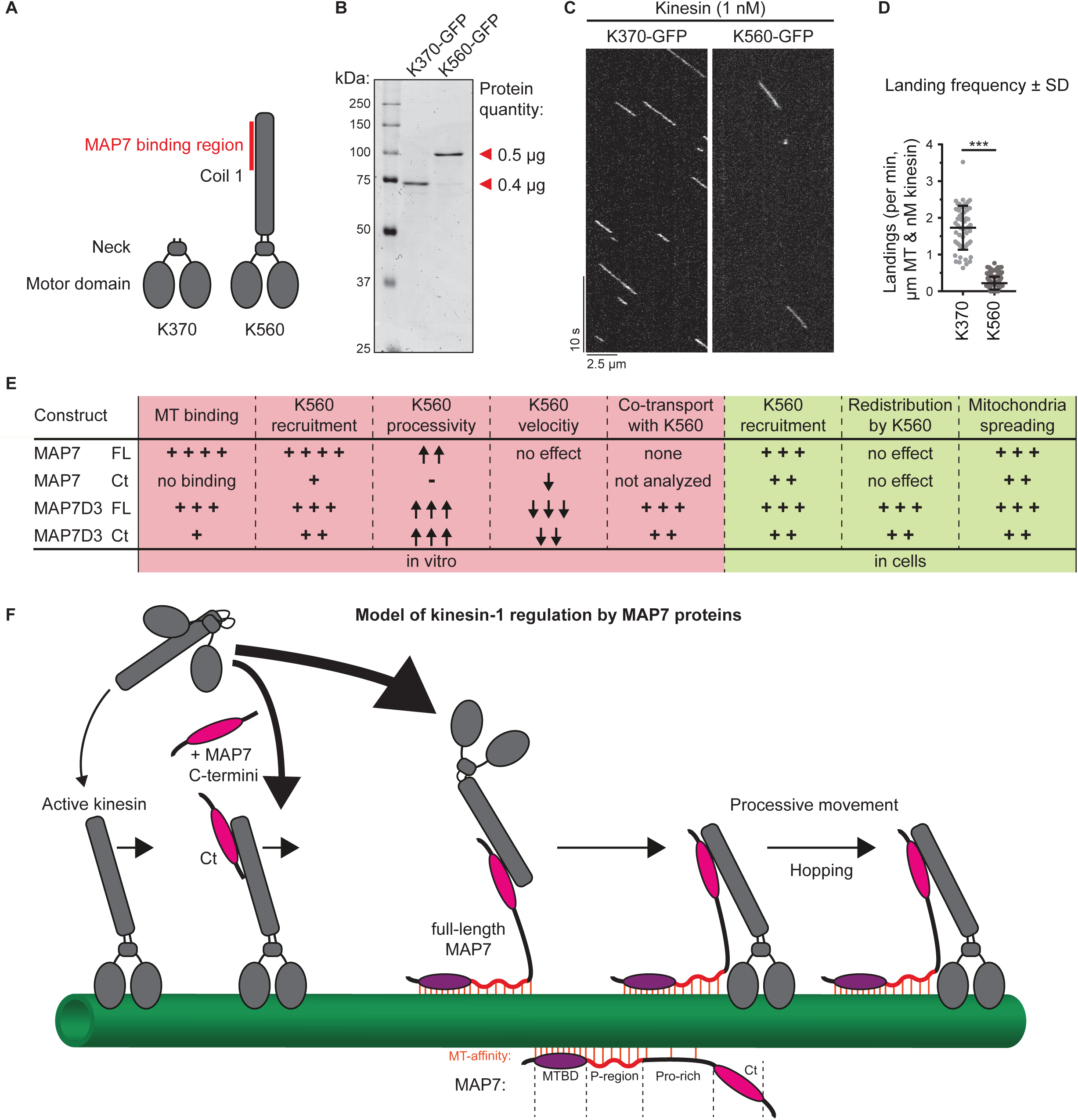
Removal of the kinesin-1 stalk domain enhances motor landing. (A) Overview of kinesin-1 constructs used. (B) Purified K370-GFP and K560-GFP analyzed by SDS-PAGE. Protein concentrations were determined using BSA standard. (C) Kymographs of K370-GFP and K560-GFP motors moving on dynamic MTs. (D) Quantification of kinesin landing frequencies, n = 57 (K370-GFP) from three independent experiments and n = 167 (K560-GFP) from two independent experiments, Mann-Whitney U test: ***, P = <0.001. (E) Summarizing table of the characteristics and effects of MAP7 proteins on different parameters of kinesin activity observed in vitro and in cells. (F) Working model of kinesin-1 activation by MAP7 family proteins. Red lines indicate MAP-MT interactions.

### Discussion

In this study, we have systematically analyzed the impact of mammalian MAP7 family proteins on kinesin-1 transport (Fig. 9E). We found that at least one MAP7 homolog was necessary and sufficient to enable kinesin-1-driven mitochondria distribution. These results are in agreement with the data showing that MAP7/ensconsin is an essential kinesin-1 co-factor in flies and in mammalian muscle cells (Barlan et al., 2013; Metivier et al., 2018; Metzger et al., 2012; Monroy et al., 2018; Sung et al., 2008). Dependence on MAP7 family members likely applies to many other kinesin-1-driven processes in mammals, because the core part of kinesin-1, the K560 fragment, was unable to bind to MTs efficiently in cells when all MAP7 homologues were absent.

Kinesin-1 function could be rescued to a significant extent by a MAP7 fragment that binds to kinesin, but has only low MT affinity, again in agreement with the data obtained in *Drosophila* (Barlan et al., 2013; Metivier et al., 2018). K560 recruitment to MTs was also stimulated by this MAP7 fragment both in cells and in vitro. This effect can likely be explained by a combination of two factors: the weak residual MT affinity present in the unstructured part of MAP7-Ct (Pro-rich region) as well as an allosteric regulation of the K560 motor by the kinesin-binding domain of MAP7. The ability of MAP7 to affect K560 allosterically is supported by the increased landing rate of the “self-activating” KIF5B-MAP7 fusion, which lacks all MAP7 linker sequences with potential MT affinity. This notion is also supported by the observation that a short kinesin-1 version (K370) lacking the MAP7-binding stalk region interacts with MTs more efficiently than K560. We propose that the stalk-containing kinesin can adopt conformations that are unfavorable for MT binding, whereas the interaction with MAP7 allosterically stabilizes a conformation that promotes landing on MTs (Fig. 9F). Diverse regulatory steps have been described for kinesin-1, mostly involving the autoinhibitory C-terminal tail region (reviewed in (Verhey and Hammond, 2009)) and also the motor domain (Xu et al., 2012). Our data add to this complexity by showing that the state of the stalk and its binding partners might directly affect kinesin interaction with MTs.

Although a MAP7 fragment lacking MT affinity can promote the engagement of kinesin-1 with MTs, the presence of MT binding regions makes this regulation much more efficient and robust. Without it, a very high concentration of the kinesin binding domain or its direct fusion to the kinesin were required to activate the kinesin. The presence of MT binding sites concentrates the kinesin binding domain of the MAP on MTs and can thus facilitate its interaction with the motor (Fig. 9F). Importantly, MT-bound MAP7 proteins have a significant effect not only on the kinesin landing rate but also on its processivity, and we excluded the possibility that this was due to kinesin multimerization. Interestingly, both a very immobile MAP (MAP7) and a more dynamic and diffusively behaving MAP (MAP7D3) could enhance kinesin-1 processivity, but the latter was able to exert this effect when present on a MT at a lower density. An obvious hypothesis is that kinesin processivity is governed by the presence of additional links to MTs. However, the distribution of run lengths of K560 was monoexponential even in the presence of MAP7 proteins, whereas a significant contribution of an additional MAP-dependent MT bound state would be expected to lead to a distribution corresponding to the sum of two or three exponential decays (Klumpp and Lipowsky, 2005). These data suggest that the interaction with MT-bound MAP7’s does not create an additional connection to a MT but somehow alters the conformational state of the motor to prevent its dissociation from MTs. Additional work would be needed to understand why a MT-unbound MAP7 fragment is sufficient to promote kinesin-1 landing but does not increase its processivity even when directly fused to the motor, whereas a MT-bound MAP7 version can stimulate not only the initial kinesin engagement with a MT but also its processive motility.

The relatively low affinity of the MAP-kinesin interaction and the rapid binding-unbinding kinetics suggest that the kinesin could be “hopping” from one MAP molecule to another, and this can allow even a highly immobile MAP7 to counteract kinesin dissociation without strongly affecting motor velocity. MAP7D3 is different from MAP7 because it binds less tightly to MTs and more tightly to the kinesin and can be “dragged” with the motor to some extent. This possibly explains why it slows down kinesin movement and why its low concentration is sufficient to increase motor processivity. However, because of the fast turnover within the MAP-motor complex, MAP7D3 is unlikely to be undergoing large-distance transport by the kinesin in cells. MAP7D3 can be rapidly relocalized to MT plus ends by overexpressed K560, but at the endogenous kinesin-1 expression levels, MAP7D3 is not enriched at MT plus ends, suggesting that the levels of endogenous motor are insufficient to drive MAP7D3 to the cell periphery. Still, it is possible that also the endogenous kinesin-1 could cause some redistribution of its own positive regulators, which would be in line with some previous observations (Kikuchi et al., 2018).

An important question is whether the localization of MAP7 proteins contributes to the well-documented selectivity of kinesin-1 for specific MT tracks, which appear to correspond to the stable, long-lived MT populations (Cai et al., 2009; Farias et al., 2015; Guardia et al., 2016; Hammond et al., 2008; Jacobson et al., 2006; Nakata and Hirokawa, 2003; Tas et al., 2017). MAP7, which is stably associated with MTs, could potentially predispose kinesin-1 for interacting with more long-lived MTs, on which this MAP would gradually accumulate. It would be interesting to know if this is indeed the case and whether different combinations of MAP7 family proteins with different properties provide fine-tuning of kinesin-1-dependent processes, particularly in complex cell types such as neurons. Taken together, our data illustrate the complexity of the interplay between the motors and the tracks they use during intracellular transport processes.

## Materials and Methods

### Cell Culture, knockdowns and CRISPR/Cas9 knockouts

HeLa (Kyoto), COS7 and human embryonic kidney 239T (HEK293T) cell lines were cultured in medium that consisted of 45% DMEM, 45% Ham’s F10, and 10% fetal calf serum supplemented with penicillin and streptomycin. The cell lines were routinely checked for mycoplasma contamination using LT07-518 Mycoalert assay (Lonza). HeLa and COS7 cells were transfected with plasmids using FuGENE 6 (Promega) for generating knockout lines, live cell imaging and immunofluorescence experiments. For streptavidin pull down assays and protein purification from HEK293T cells, plasmids were transfected with polyethylenimine (PEI; Polysciences). For generating knockdowns, HeLa cells were transfected with 100 nM siRNA for each target using HiPerfect (Qiagen). The following siRNAs were used in this study: MAP7D1 (target sequence 5’-TCATGAAGAGGACTCGGAA-3’), MAP7D3 (target sequence 5’-AACCTACATTCGTCTACTGAT-3’) and luciferase (target sequence 5’-TCGAAGTATTCCGCGTACG-3’). For rescue experiments, HeLa cells were transfected 2 days after siRNA transfection.

HeLa CRISPR/Cas9 knockout lines were generated using the pSpCas9-2A-Puro (PX459) vector, purchased from Addgene (Ran et al., 2013). Guide RNAs for human KIF5B, MAP7, and MAP7D3 were designed using the CRISPR design webpage tool (http://crispr.mit.edu). The targeting sequences for gRNAs were as follows (coding strand sequence indicated): KIF5B, 5′-CCGATCAAATGCATAAGGCT-3′; MAP7, 5′-CGCCCTGCCTCTGCAATTTC-3′; MAP7D3, 5′-CCGTGCCCGCAGCTCTCTCA-3′. The CRISPR/Cas9-mediated knockout of KIF5B, MAP7, and MAP7D3-encoding genes was performed according to the protocol described in (Ran et al., 2013). In brief, HeLa cells were transfected using Fugene6 (Promega) with the vectors bearing the appropriate targeting sequences. Cells were subjected to selection with 2 µg/mL puromycin 24-48 hours post transfection for 48-72 hours. After selection, cells were allowed to recover in complete medium for approximately 7 days; meanwhile, cells were diluted in 96 wells plates for growing single cell colonies. Knockout efficiency was controlled using mixed cell populations by immunofluorescence staining, and depending on the efficiency, 5-30 individual clones were isolated and characterized by Western blotting and immunostaining.

### DNA constructs

All MAP7 family protein constructs except MAP7D1-FL were cloned by a PCR-based strategy into a pBio-mCherry-C1 vector, a modified pEGFP-C1 vector, in which the open reading frame encoding EGFP was substituted for mCherry, and a linker encoding the sequence MASGLNDIFEAQKIEWHEGGG, which is the substrate of biotin ligase BirA, was inserted into the NheI and AgeI sites in front of the mCherry-encoding sequence. MAP7D1-FL has been cloned into a Bio-mCherry-C3 vector in a similar fashion. MAP7 constructs were generated using cDNA of RNA isolated from HeLa cells. MAP7D1 was derived from the IMAGE clone 6514558 (Source Bioscience), MAP7D2 was derived from the IMAGE clone 3063898 (Source Bioscience) and MAP7D3 was from the IMAGE clone 5284128 (Source Bioscience). All MAP7 linker constructs were cloned in a pEGFP-C1 vector by a PCR-based strategy. Rescue constructs for MAP7D1 and MAP7D3 were obtained by PCR-based mutagenesis of the sequence TCATGAAGAGGACTCGGAA to TCATGAAGAGAACACGCAA (MAP7D1) and AACCTACATTCGTCTACTGAT to AATCTACACTCGTCTACAGAT (MAP7D3). For protein purification from HEK293T cells, MAP7-Ct was cloned into a pmCherry-C1 vector with an N-terminal Strep-tag, and full-length MAP7 and MAP7D3 were cloned into a pTT5 vector with an N-terminal SNAP- and Strep-tag. For bacterial purifications, MAP7D3 constructs were cloned into a pET28a vector containing an N-terminal Strep-tag. MAP7-Ct(mini) construct was cloned by a PCR-based strategy into a pET24a vector containing an N-terminal SUMO-tag. All KIF5B constructs were cloned by a PCR-based strategy into pEGFP-N1 vectors. K560-GFP and full-length KIF5B-GFP used for protein purification were cloned into a pTT5 vector with a C-terminal Strep-tag. K370-GFP, saK560-GFP and saK560-GFP(Δ) used for protein purification and cellular experiments were cloned into a pEGFP-N1 vector with a C-terminal Strep-tag. KIF5B-Coil2-GFP and KIF5B-Tail-GFP were cloned in an pEGFP-C1 vector. All kinesin constructs were based on full length human KIF5B as template IMAGE clone 8991997 (van Spronsen et al., 2013). KIF5B-mCherry-LexyC was cloned by PCR-based strategy. A c-myc NLS sequence was introduced between K560 and mCherry (amino acid sequence: PAAKRVKLD) and a second SV40 Large T-antigen nuclear localization signal (NLS) was introduced between mCherry and the LEXY domain (amino acid sequence: PKKKRKV). The engineered LOV2-domain (LEXY) for blue light-inducible nuclear export was obtained from Addgene (catalog number #72655) (Niopek et al., 2016). For affinity measurements, minimal protein fragments for MAP7 and MAP7D3 where designed based on structure prediction and intron-exon analysis. For KIF5B, the designed fragment was the shortest construct still containing the cysteine at position 421 and the MAP7 binding domain (aa 508-523) (Monroy et al., 2018). All protein constructs were fused to an N-terminal thioredoxin-6xHis cleavable tag by restriction free positive selection into a pET-based bacterial expression vector by a PCR-based strategy (Olieric et al., 2010). Biotin ligase BirA expression construct (Driegen et al., 2005) was a kind gift from D. Meijer (University of Edinburgh, UK).

### Antibodies, Western blotting and immunofluorescence cell staining

For immunofluorescence cell staining and Western blotting, we used rabbit polyclonal antibodies against MAP7D1 (HPA028075, Sigma/Atlas), MAP7D2 (HPA051508, Sigma/Atlas), MAP7D3 (HPA035598, Sigma/Atlas), Kinesin heavy chain (UKHC H-50, SC28538, Santa Cruz) and GFP (ab290; Abcam). We used a mouse polyclonal antibody against MAP7 (H00009053-B01P, Abnova) and mouse monoclonal antibodies against Ku80 (611360, BD Bioscience), mCherry (632543, Clontech), Cytochrome c (556432, BD Bioscience) and α-tubulin (T6199, Sigma), and a rat monoclonal antibody against α-tubulin (γL1/2, ab6160, Abcam). The following secondary antibodies were used: IRDye 800CW/680LT goat anti–rabbit, and anti–mouse for Western blotting and Alexa Fluor 488–, 594–, and 647–conjugated goat antibodies against rabbit, rat, and mouse IgG (Molecular Probes) for immunofluorescence. Mitotracker Red CMXRos (Molecular Probes) was alternatively used for mitochondria staining.

Total HeLa cell extracts were prepared in RIPA buffer containing 50 mM Tris-HCl pH 7.5, 150 mM NaCl, 1% Triton X-100, 0,5% SDS and cOmplete protease inhibitor cocktail (Roche).

For immunofluorescence cell staining, HeLa cells were fixed in –20°C methanol for 10 min and stained for MAP7, MAP7D1, MAP7D2, MAP7D3 and α-tubulin. In the case of cytochrome c, cells were fixed with 4% PFA in phosphate-buffered saline (PBS) for 10 min. Cells were then permeabilized with 0.15% Triton X-100 in PBS for 2 min; subsequent wash steps were performed in PBS supplemented with 0.05% Tween-20. Epitope blocking and antibody labeling steps were performed in PBS supplemented with 0.05% Tween-20 and 1% BSA. Before mounting in Vectashield mounting medium (Vector Laboratories) slides were washed with 70% and 100% ethanol and air-dried.

### Pull down assays

Streptavidin pull down assays were performed from HEK293T cell lysates by coexpressing biotin ligase BirA with mCherry-tagged constructs containing a biotinylation site (pBio-mCherry) (bait), and GFP-tagged KIF5B constructs (prey). Constructs were transfected altogether into HEK293 cells using PEI with 24 hrs incubation time for protein expression. M-280 Streptavidin Dynabeads (Invitrogen) were blocked in a buffer containing 20 mM Tris pH 7.5, 20% glycerol, 150 mM NaCl, and 10 µg Chicken Egg Albumin followed by three washes with wash buffer containing 20 mM Tris pH 7.5, 150 mM NaCl and 0.1% Triton-X. HEK293T cells were scraped and collected in ice-cold PBS followed by lysis on ice in a buffer containing 20 mM Tris pH 7.5, 150 mM NaCl, 1mM MgCl_2_, 1% Triton X-100, and cOmplete protease inhibitor cocktail (Roche). To separate cell debris, the lysates were cleared by centrifugation at 4°C for 15 min at 16000 g and 10 % of each lysate was saved as input control. Cell lysates were incubated with pre-blocked streptavidin beads for 60 min at 4°C followed by five washes with the buffer containing 20 mM Tris pH 7.5, 150 mM NaCl and 0.1% Triton-X. Streptavidin beads were pelleted and boiled in 2x Laemmli sample buffer. Protein lysates and pull down of both bait and prey proteins were analyzed by Western blotting.

### Protein Purification

All KIF5B, SNAP(Alexa647)-labeled proteins and mCherry-MAP7-Ct used for in vitro reconstitution assays were purified from HEK293T cells using Strep(II)-streptactin affinity purification. Cells were harvested 24-40 hrs post transfection. Cells from a 15 cm cell culture dish were lysed in 800 μL of lysis buffer containing 50 mM HEPES pH 7.4, 300 mM NaCl, 1 mM MgCl_2_, 1 mM DTT and 0.5% Triton X-100, supplemented with cOmplete protease inhibitor cocktail (Roche) on ice for 15 min. The supernatants obtained from cell lysates after centrifugation at 16000 x g for 20 min were incubated with 50 μL of StrepTactin Sepharose beads (GE Healthcare) for 1 hr. The beads were washed 5 times in a high-salt wash buffer containing 50 mM HEPES pH 7.4, 1.5 M NaCl, 1 mM MgCl_2_, 1 mM EGTA, 1 mM DTT, and 0.05% Triton X-100 and three times with an elution wash buffer containing containing 50 mM HEPES pH 7.4, 150 mM NaCl, 1 mM MgCl_2_, 1 mM EGTA, 1 mM DTT and 0.05% Triton X-100. The proteins were eluted with 40-150 μL of elution buffer containing 50 mM HEPES pH 7.4, 150 mM NaCl, 1 mM MgCl_2_, 1 mM EGTA, 1 mM DTT, 2.5 mM d-Desthiobiotin and 0.05% Triton X-100. To label SNAP-tagged proteins with SNAP-Surface Alexa Fluor 647 (NEB), 20– 40 μM dye was incubated with proteins on beads for 1 hr between wash and elution steps. After extensive washing, proteins were eluted in the elution buffer with 300 mM instead of 150 mM NaCl. The concentrations of purified proteins were measured by BSA standard using SDS-PAGE. All purified proteins were snap frozen in liquid nitrogen and stored in −80°C.

For protein purification of MAP7D3 constructs from bacteria, BL21 *E.coli* were transformed with the respective MAP7D3 construct. Bacteria were grown till OD600 of 0.6 at 37°C after which protein expression was induced with 1 mM IPTG for 1 hr at 37°C and 2.5 hrs at 20°C. Bacteria were spun down and subjected to one freeze-thaw cycle at −80°C to stimulate proper lysis. Bacteria were resuspended and sonicated at 4°C in cold lysis buffer containing 50 mM sodium phosphate pH 8, 250 mM NaCl, 1 mM MgCl_2_, 1 mM DTT, 0.5% Triton X-100, 1mM PMSF and cOmplete protease inhibitor cocktail (Roche). Lysates were centrifuged at 25000 x g for 45-60 min and the supernatants were incubated with Strep-Tactin Superflow high-capacity beads (IBA Lifesciences) for 60 min at 4°C, followed by 5 washes with a buffer containing 50 mM sodium phosphate pH 6.0, 250 mM NaCl, 1 mM MgCl_2_ and 1 mM DTT. The proteins of interest were eluted with a buffer containing 50 mM sodium phosphate pH 7.0, 250 mM NaCl, 1 mM MgCl_2_ 1 mM DTT and 5 mM d-Desthiotiotin. The eluted fractions were pooled and supplemented with 10% sucrose for preservation. Proteins were snap frozen and stored at −80°C. The concentrations of purified proteins were measured by BSA standard using SDS-PAGE.

For protein purification of the minimal MAP7, MAP7D3 and KIF5B fragments used for probing complex formation (SEC-MALS and ITC), all protein constructs were fused to an N-terminal thioredoxin-6xHis cleavable tag by restriction free positive selection into a pET-based bacterial expression vector. For expression, the, KIF5B and MAP7D3 constructs were transformed using *E. coli* expression strain BL21(DE3), whereas for MAP7 construct BL21-CodonPlus (DE3)-RIPL competent cells were chosen. All transformed cells were cultivated in LB containing 50 µg/mL Kanamycin. Subsequently, upon reaching an OD600 of 0.4 to 0.6 the cultures were induced with 1 mM isopropyl 1-thio-β-galactopyranoside (IPTG, Sigma) after cooling to 20°C for KIF5B and MAP7D3 or at 37°C for MAP7 respectively. The KIF5B and MAP7D3 constructs were expressed at 20°C overnight whereas the MAP7 construct was expressed for 4 h at 37°C. Cells were harvested by centrifugation at 4000 x g at 4°C followed by sonication for lysis in the buffer containing 50 mM HEPES, pH 7.5, 500 mM NaCl, 10 mM Imidazole, 10% Glycerol, 2 mM β-mercaptoethanol, proteases inhibitors (Roche) and DNAseI (Sigma). The lysate was cleared by centrifugation at 18000 x g for 20 min and subsequent filtration through a 0.45 µm filter. All constructs were purified by immobilized metal-affinity chromatography (IMAC) on HisTrap HP Ni^2+^ Sepharose columns (GE Healthcare) at 4°C according to the manufacturer's instructions. The elutions were cleaved by 3C protease during dialysis against the same buffer as above lacking protease inhibitors and DNAse I. The cleaved constructs were separated from tag and protease by a reverse IMAC run followed by a gel filtration run on a HiLoad Superdex 200 16/60 size exclusion chromatography column (GE Healthcare) in the buffer containing 10 mM Tris HCl, pH 7.5 and 150 mM NaCl. Fractions containing the constructs were pooled and concentrated by ultracentrifugation before being flash frozen and stored at −80°C.

For recombinant protein production of MAP7-Ct(mini), BL21(DE3) Rosetta2 *E. coli* cells (Novagen) containing a pET24a vector (Novagen) encoding a SUMO-MAP7-Ct(mini) construct were cultured. 2 L culture, in LB medium supplemented with the antibiotics 10 g/L kanamycin (Sigma-Aldrich) and 33 g/L chloramphenicol (Sigma-Aldrich), were grown until OD600 of ~0.8-1, after which protein production was induced with 0.1 mM IPTG (Thermo Scientific). Protein production was performed overnight at 18°C. Cells were harvested by centrifugation and subjected to one freeze-thaw cycle at −80°C to initiate cell lysis. The pellet was thawed and resuspended in 50 mM sodium phosphate buffer pH 8.0, 150 mM NaCl, cOmplete protease inhibitor cocktail (Roche) and 5 mM β-mercaptoethanol. The cells were then disrupted by an EmulsiFlex-C5 (Avestin) cell disruptor. The lysate was cleared by centrifugation (55,000 g, 45 min), filtered through a 0.22 μm polypropylene filter (VWR) and mixed for 15 min with Protino^®^ Ni-IDA resin (Macherey-Nagel) at 4°C. After centrifugation, the protein was eluted with 50 mM sodium phosphate pH 8.0, 250 mM imidazole, 150 mM NaCl, cOmplete protease inhibitor cocktail (Roche) and 5 mM β-mercaptoethanol. To cleave off the SUMO tag, the eluate was then digested with Ulp1 overnight at 4°C while dialyzed against 50 mM phosphate buffer, pH 8.0 with a 6 kDa cut-off membrane (Spectrum Laboratories). The protein was loaded on a POROS® 20HS (Thermo Fischer Scientific) cation exchange column in the same dialysis buffer, using an ÄKTA® Purifier chromatography system (GE Healthcare). The protein was eluted by a linear gradient up to 2 M KCl over 15 CV (Carl Roth); fractions of 0.5 mL were collected. Fractions of interest were then concentrated and exchanged against 25 mM HEPES buffer pH 7.5 with 75 mM KCl, 75 mM NaCl and 10 mM DTT using a Vivaspin column (cut-off: 6 kDa). The final concentration was determined by an ND-100 spectrophotometer (Nanodrop Technologies). Purity was confirmed by SDS-PAGE and the protein was aliquoted and stored at −80°C.

### Mass spectrometry

After streptavidin purification, beads were resuspended in 20 µl of Laemmli sample buffer (Biorad) and supernatants were loaded on a 4-12% gradient Criterion XT Bis-Tris precast gel (Biorad). The gel was fixed with 40% methanol/10% acetic acid and then stained for 1 hr using colloidal Coomassie dye G-250 (Gel Code Blue Stain Reagent, Thermo Scientific). After in-gel digestion, samples were resuspended in 10% formic acid (FA)/5% DMSO and analyzed using an Agilent 1290 Infinity (Agilent Technologies, CA) LC, operating in reverse-phase (C18) mode, coupled to an Orbitrap Q-Exactive mass spectrometer (Thermo Fisher Scientific, Bremen, Germany). Peptides were loaded onto a trap column (Reprosil C18, 3 µm, 2 cm × 100 µm; Dr. Maisch) with solvent A (0.1% formic acid in water) at a maximum pressure of 800 bar and chromatographically separated over the analytical column (Zorbax SB-C18, 1.8 µm, 40 cm × 50 µm; Agilent) using 90 min linear gradient from 7-30% solvent B (0.1% formic acid in acetonitrile) at a flow rate of 150 nL/min. The mass spectrometer was used in a data-dependent mode, which automatically switched between MS and MS/MS. After a survey scan from 350-1500 m/z the 10 most abundant peptides were subjected to HCD fragmentation. MS spectra were acquired in high-resolution mode (R > 30,000), whereas MS2 was in high-sensitivity mode (R > 15,000). Raw files were processed using Proteome Discoverer 1.4 (version 1.4.0.288, Thermo Scientific, Bremen, Germany). The database search was performed using Mascot (version 2.4.1, Matrix Science, UK) against a Swiss-Prot database (taxonomy human). Carbamidomethylation of cysteines was set as a fixed modification and oxidation of methionine was set as a variable modification. Trypsin was specified as enzyme and up to two miss cleavages were allowed. Data filtering was performed using percolator, resulting in 1% false discovery rate (FDR). Additional filters were; search engine rank 1 peptides and ion score >20.

### Size exclusion chromatography followed by multi-angle light scattering (SEC-MALS) experiments

SEC-MALS experiments were conducted in a buffer containing 10 mM Tris HCl, pH 7.5 supplemented with 150 mM NaCl at 20°C using a Superdex 200 10/300 analytical size exclusion chromatography column (GE Healthcare) coupled to a miniDAWN TREOS light scattering and Optilab T-rEX refractive index detector (Wyatt Technology). A volume of 30 µL of protein samples at 5 mg/mL was injected and data was analyzed with the software provided by the manufacturer.

### Isothermal titration calorimetry (ITC)

The proteins were dialyzed against a buffer containing 10 mM Tris HCl, pH 7.5, supplemented with 150 mM NaCl overnight at 4°C. ITC experiments were conducted on an iTC200 instrument at 25°C, using an injection volume of 2.1 µL at a reference power of 7 and stirring speed of 700 rpm. For probing the MAP7-KIF5B interaction, the MAP7 fragment was used as titrant at a concentration 576 µM against the KIF5B fragment at 15 µM in the cell. The MAP7D3-KIF5B interaction was probed with 300 µM MAP7D3 fragment as titrant and 30 µM KIF5B fragment in the cell. The resulting enthalpograms were integrated and fitted using the standard one-site-model of Origin (OriginLab).

### In vitro reconstitution assays

MT seeds were prepared by incubating 20 μM porcine tubulin mix containing 70% unlabeled, 18% biotin-tubulin and 12% rhodamine-tubulin with 1 mM guanylyl-(α,β)-methylenediphosphonate (GMPCPP) at 37°C for 30 min. Polymerized MTs were separated from the mix by centrifugation in an Airfuge at 119,000 g for 5 min. MTs were subjected to one round of depolymerization and polymerization in 1 mM GMPCPP, and the final MT seeds were stored in MRB80 buffer (80 mM K-PIPES pH 6.8, 1 mM EGTA, 4 mM MgCl_2_) containing 10% glycerol. In vitro reconstitution assays were performed in flow chambers assembled from microscopy slides and plasma cleaned coverslips. The chambers were treated with 0.2 mg/mL PLL-PEG-biotin (Surface Solutions, Switzerland) in MRB80 buffer for 5 min. After washing with the assay buffer, they were incubated with 1 mg/mL NeutrAvidin for 5 min. MT seeds were attached to the biotin-NeutrAvidin links and incubated with 1 mg/mL κ-casein. The in vitro reaction mixture consisted of 20 μM tubulin, 50 mM KCl, 0.1% methylcellulose, 0.5 mg/mL κ-casein, 1 mM GTP, oxygen scavenging system (20 mM glucose, 200 μg/mL catalase, 400 μg/mL glucose-oxidase, 4 mM DTT), 2 mM ATP, 0.2 - 10 nM of respective kinesin (concentrations were calculated for monomeric proteins) and MAP7 or MAP7D3 at indicated concentrations. After centrifugation in an Airfuge for 5 min at 119,000 g, the reaction mixture was added to the flow chamber containing the MT seeds and sealed with vacuum grease. The experiments were conducted at 30°C and data were collected using TIRF microscopy. For some experiments without mCherry-labeled MAP7/MAP7D3, the reaction mixture was composed of 19.5 μM tubulin supplemented with 0.5 μM rhodamine-labeleld tubulin. All tubulin products were purchased from Cytoskeleton Inc.

For MT pelleting assays a reaction containing 37.5 μM porcine brain tubulin supplemented with 1 mM GTP, 1 mM DTT and 20 μM Taxol in MRB80 was prepared at 30°C for 30 min. The reaction was divided and supplemented with MRB80 buffer with or without mCherry-MAP7-Ct at a final tubulin concentration of 30 μM. Also a control without tubulin was included. Subsequently, all reactions were incubated for another 15 min at 30°C. Pelleting was performed in an Airfuge at 119000 g with a pre-warmed rotor for 10 min. Supernatants were removed and pellets were resuspended in MRB80 buffer on ice by regular pipetting for 40 min. All samples were supplemented with 4x Laemmli sample buffer, boiled and analyzed by SDS-PAGE.

### Image acquisition

Fixed cells were imaged with a Nikon Eclipse 80i upright fluorescence microscope equipped with Plan Apo VC N.A. 1.40 oil 100x and 60x objectives, or Nikon Eclipse Ni-E upright fluorescence microscope equipped with Plan Apo Lambda 100x N.A. 1.45 oil and 60x N.A. 1.40 oil objectives microscopes, Chroma ET-BFP2, - GFP, -mCherry, or -Cy5 filters and Photometrics CoolSNAP HQ2 CCD (Roper Scientific, Trenton, NJ) camera. The microscopes were controlled by Nikon NIS Br software.

FRAP and LEXY optogenetic experiments were done using spinning disk microscopy, which was performed on an inverted research microscope Eclipse Ti-E with the Perfect Focus System (Nikon), equipped with Plan Apo VC 100x N.A. 1.40 and Plan Apo 60x N.A. 1.40 oil objectives, a Yokogawa CSU-X1-A1 confocal head with 405-491-561 triple-band mirror and GFP, mCherry, and GFP/mCherry emission filters (Chroma), ASI motorized stage MS-2000-XYZ with Piezo Top Plate (ASI), a Photometrics Evolve 512 electron-multiplying charge-coupled device (CCD) camera (Photometrics), and controlled by MetaMorph 7.7 software (Molecular Devices). The microscope was equipped with a custom-ordered illuminator (MEY10021; Nikon) modified by Roper Scientific France/PICT-IBiSA, Institut Curie. Cobolt Calypso 491 nm (100 mW) and Cobolt Jive 561 nm (100 mW) lasers (Cobolt) were used as light sources. To keep cells at 37°C, we used a stage top incubator (model INUBG2E-ZILCS; Tokai Hit).

All in vitro reconstitution assays were imaged on an inverted research microscope Nikon Eclipse Ti-E (Nikon) with the perfect focus system (PFS) (Nikon), equipped with Nikon CFI Apo TIRF 100 × 1.49 N.A. oil objective (Nikon, Tokyo, Japan), Photometrics Evolve 512 EMCCD (Roper Scientific) and Photometrics CoolSNAP HQ2 CCD (Roper Scientific) and controlled with MetaMorph 7.7 software (Molecular Devices, CA). The microscope was equipped with TIRF-E motorized TIRF illuminator modified by Roper Scientific France/PICT-IBiSA, Institut Curie. For excitation lasers we used 491 nm 100 mW Stradus (Vortran), 561 nm 100 mW Jive (Cobolt) and 642 nm 110 mW Stradus (Vortran). We used an ET-GFP 49002 filter set (Chroma) for imaging of proteins tagged with GFP, an ET-mCherry 49008 filter set (Chroma) for imaging X-Rhodamine labelled tubulin or mCherry-tagged proteins and an ET-405/488/561/647 for imaging SNAP-Alexa647. For simultaneous imaging of green and red fluorescence we used an Evolve512 EMCCD camera (Photometrics), ET-GFP/mCherry filter cube (59022, Chroma) together with an Optosplit III beamsplitter (Cairn Research Ltd) equipped with double emission filter cube configured with ET525/50m, ET630/75m and T585lprx (Chroma). For simultaneous imaging of green, red and far-red fluorescence we used an Evolve512 EMCCD camera (Photometrics), quad TIRF polychroic ZT405/488/561/640rpc (Chroma) and quad laser emission filter ZET405/488/561/635m (Chroma), mounted in the metal cube (Chroma, 91032) together with an Optosplit III beamsplitter (Cairn Research Ltd) equipped with triple emission filter cube configured with ET525/50m, ET630/75m, ET700/75m emission filters and T585lprx and T660lprx dichroic (Chroma). To keep in vitro samples at 30°C, we used a stage top incubator (model INUBG2E-ZILCS; Tokai Hit).

### Image processing and analysis

Images and movies were processed using ImageJ. All images were modified by linear adjustments of brightness and contrast. Maximum intensity projections were made using z projection. Kinesin velocities, run lengths and landing frequencies were obtained from kymograph analysis, using ImageJ plugin KymoResliceWide v.0.4. https://github.com/ekatrukha/KymoResliceWide; copy archived at https://github.com/elifesciences-publications/KymoResliceWide). Kinesin runs <0.5 sec were included for landing frequency analysis but not analyzed for run length and velocity. Kinesins running on GMPCPP MT seeds were excluded from our analysis as much as possible. Kinesin runs >2.0 sec were analyzed for MAP7/MAP7D3 co-transport events.

For quantifying mitochondria in different HeLa cell knockdown/knockout conditions, we classified mitochondria as “clustered” when ~80% of the cytochrome c signal was localized in a dense cluster around the nucleus, whereas all other localization patterns with more spread mitochondria were classified as “spread”. K560-GFP was classified as localized on corner MTs when clear enrichment of fluorescent signal was seen at MT ends near the cell periphery over MTs that are localized in between the boundary and the cell center.

Line scan analysis of MT localization of MAP7 linker truncations was performed on widefield images of HeLa and COS7 cells overexpressing GFP-tagged constructs co-stained for α-tubulin. Average fluorescence intensities were measured from 1-3 micron line scans along MTs stained for α-tubulin and an adjacent line 5 pixels away from the same MT as a background intensity measurement. Next, the ratio was calculated by dividing fluorescent intensity of the MT line scan by the fluorescent intensity of the control line scan at the same region. No enrichment of the GFP-tagged construct on MTs would result in an intensity ratio of 1. Per condition, 30 MTs were analyzed form 10 cells (3 MTs per cell).

Imaging of MT plus ends with EB3-GFP as a marker was performed on a TIRF microscope. Imaging was performed at 2 frames per second for 50 seconds. Per cell, approximately 10-30 EB3 comets were traced and average growth velocity/duration was calculated and analyzed per cell using SigmaPlot.

FRAP measurements were performed by bleaching a 10 × 10 μm square region in a cytoplasmic region between the nucleus and cell cortex followed by 8.5 min imaging with a frame interval of 3 sec. Mean fluorescence intensities were measured from a 4 × 4 μm square region within the original photobleached region to avoid analyzing non-bleached MTs that could slide into the analyzed region. The mean intensity of this region was double corrected for background fluorescence and photobleaching (Phair et al., 2004).

Optogenetics experiments with blue light-inducible K560-LEXY kinesin were performed using spinning disk microscopy. Acquisitions were done with a frame interval of 5 sec after sequential exposure with green light 561 nm laser (to image kinesin) followed by blue light 491 nm laser (to image MAPs and activate K560-LEXY simultaneously). Exposure times of ~ 1 sec per interval with the 491 nm laser were sufficient to induce active export of LEXY-tagged motors from the nucleus. For measuring fluorescence intensity changes at cell corners, a maximum intensity projection of the K560-LEXY channel over time was made using ImageJ, followed by Gaussian blurring and thresholding to select cell corners to analyze. Mean fluorescence values for GFP-MAP7/MAP7D3 and K560-LEXY were obtained from the same cell corners over time, background subtracted and normalized to the mean fluorescence in that region at t = 0 min. Changes in mean fluorescence intensity were plotted per cell corner.

### Single molecule intensity analysis

Single molecule fluorescence histograms of monomeric GFP (control) or kinesins moving on MT lattices were built from acquisitions made on a TIRF microscope. To ensure identical imaging conditions, a single imaging slide (with a plasma cleaned coverslip) was used containing two or three flow chambers to image GFP (control) and K560-GFP (with or without MAP7 proteins) (Fig. S4C) or K370-GFP and K560-GFP (Fig. S5D). For purified GFP, the protein was diluted in MRB80 and added to an imaging flow chambers; chambers were subsequently washed with MRB80, leaving a fraction of the GFP proteins immobilized on the coverslip. Protein dilution was optimized to provide images of approximately 0.01 fluorophores per µm^2^ for GFP control conditions. To estimate the number of GFP molecules per kinesin, an in vitro reconstitution assay with K370-GFP or K560-GFP moving on MTs in the presence or absence of MAP7 proteins was set up in the other flow chambers as described before. After sealing with vacuum grease to prevent evaporation, samples were imaged at 30ºC. For monomeric GFP, approximately 100 images were acquired at different positions on the coverslip to avoid pre-bleaching. For moving kinesins, approximately 5-10 movies were obtained where only the first 10 frames were analyzed to prevent analyzing partially photobleached motors. All acquisitions were obtained under identical laser power, exposure time and TIRF angle. ImageJ plugin Comdet v.0.3.6.1 and DoM_Utrecht v.1.1.5 https://github.com/ekatrukha/DoM_Utrecht) were used for detection and fitting of single molecule fluorescent spots as described previously (Yau et al., 2014). In short, individual spots were fitted with 2D Gaussian, and the amplitude of the fitted Gaussian function was used as a measure of the fluorescence intensity value of an individual spot. The histograms were fitted to lognormal distributions using GraphPad Prism 7.

### Statistical analysis

Statistical significance was analyzed either using the Mann-Whitney U test or Student’s t test, as indicated in figure legends. For the t test, data distribution was assumed to be normal, but this was not formally tested. Statistical significance was determined using GraphPad Prism software (version 7.04). Fitting of run lengths with the sum of two or three exponential decays was performed on the raw data using maximum likelihood estimation method implemented in *mle* function of MATLAB R2011b (The MathWorks, Natick, 2011).

### Online supplemental material

Fig. S1 shows the analysis of MT organization and dynamics in MAP7 depleted cells, the distribution of MAP7D3-Ct in HeLa cells, uncropped images of the MT pelleting assay from Figure 2B and in vitro assays with mCherry-MAP7-Ct. Fig. S2 illustrates the overexpression of GFP-tagged MAP7 linker constructs in COS7 cells and overexpressed Nt and Ct constructs of MAP7, MAP7D1 and MAP7D3 together with K560-GFP in HeLa cells depleted of all three MAP7 proteins. Fig. S3 provides an overview of all purified proteins used in this study, the analysis of kinesin purification contaminants and velocity parameters of full-length KIF5B. Fig. S4 and S5 include additional control conditions for the ITC experiments, illustrate binding of MAP7 proteins to MTs in in vitro reconstitution assays as well as quantifications of kinesin behavior from Fig. 5-9. Supplemental videos show that MAP7D3 but not MAP7 can be rapidly relocalized by K560 to the cell periphery.

## Acknowledgments

We thank A. Aher for the help on protein purifications and D. Meijer for the gift of materials. This work was supported by the European Research Council Synergy grant 609822 and Netherlands Organization for Scientific Research ALW Open Program grant 824.15.017 to A.A., the Marie Curie IEF fellowships to M.M. and by a grant from the Swiss National Science Foundation (31003A_166608; to M.O.S.).

The authors declare no competing financial interests.

## Author Contributions

Author contributions: P.J. H. and M. M. designed, conducted and analyzed the experiments and wrote the paper. T.M. designed, conducted and analyzed biochemical experiments. G-J.K., C.A.E.P., D.G.F.V., W,E.vR. and I.G. contributed to cellular experiments. E.A.K. contributed to data analysis. L.F. and S.G.D.R. contributed to protein purifications. A.F.M.A. and R.S. performed and analyzed mass spectrometry experiments. C.C.H., M.O.S. and L.C.K. contributed to experiment planning, data analysis and paper writing; A. A. supervised the study and wrote the paper.

## Abbreviations

Ct: C-terminus
MAP: microtubule-associated protein
MT: microtubule
MW: molecular weight
Nt: N-terminus

## Supplemental Figure Legends

**Figure S1.**
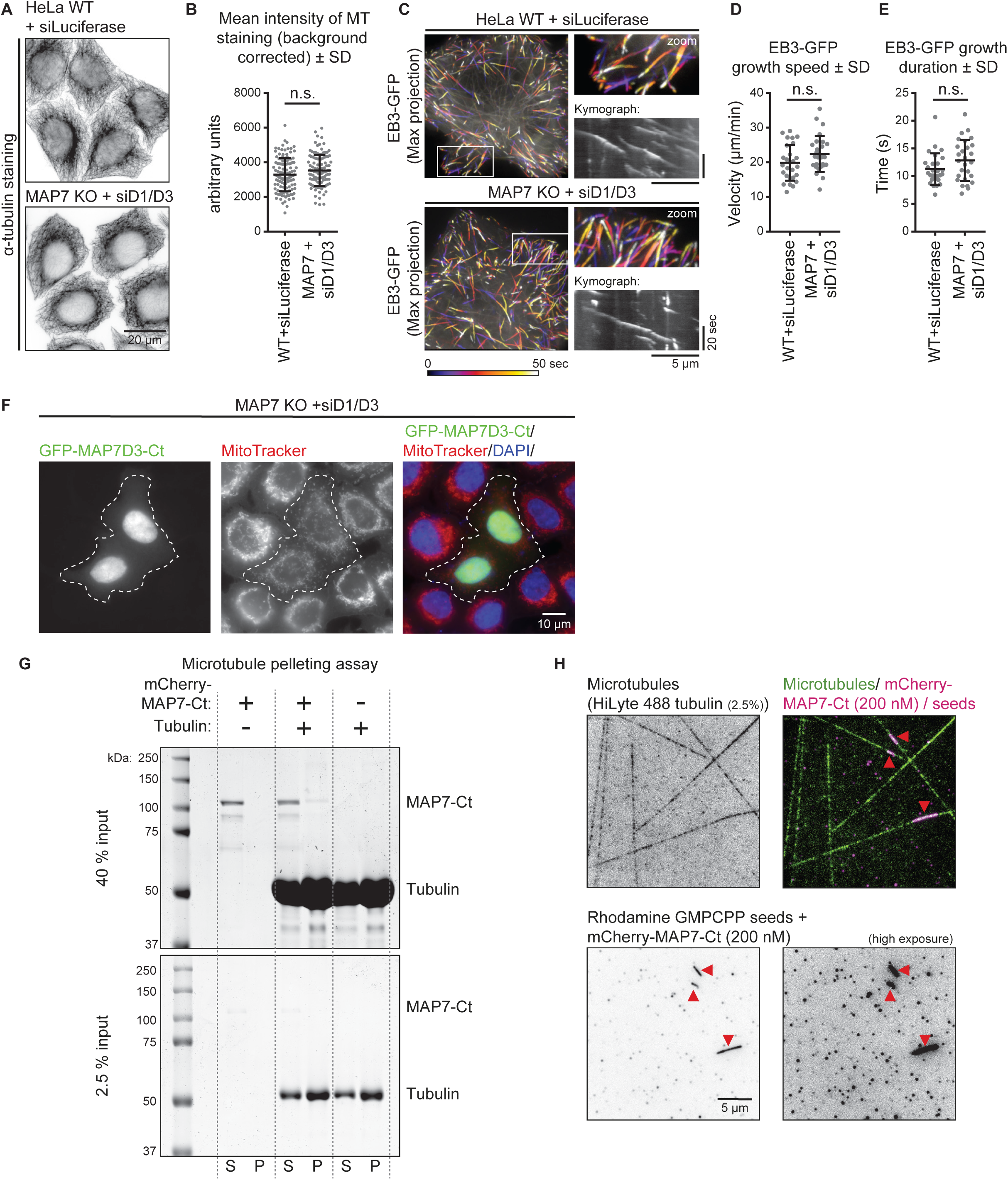
Effects of the depletion MAP7 family proteins on the MT network and characterization of their MT binding domains. (A-B) Control or MAP7 KO + siMAP7D1/D3 HeLa cells were stained for α-tubulin and imaged on a wide field microscope (A) to quantify MT intensity per cell (B); n = 123 cells (WT + siLuciferase) and n = 115 cells (MAP7 KO + siMAP7D1/D3) from three independent experiments, Student’s t-test: P = 0.054. (C) Live cell imaging of EB3-GFP in control or MAP7 KO + siMAP7D1/D3 Hela cells on a TIRF microscope. Color-coded maximum intensity projections, zooms of the white boxed area and illustrative kymographs of growing EB3-GFP comets are shown per condition. (D, E) Quantification of EB3-GFP dynamics: growth rate (D) and growth duration (E). n = 358 comets from 27 cells (WT + siLuciferase) and n = 289 comets from 27 cells (MAP7 KO + siMAP7D1/D3) from three independent experiments, Mann-Whitney U test: (D) p = 0.078 and (E) p = 0.138. (F) Wide field images of overexpressed GFP-MAP7D3-Ct in MAP7 KO + siMAP7D1/D3 HeLa cells stained with MitoTracker and DAPI to visualize mitochondria and nuclei. (G) Unprocessed Coomassie blue-stained SDS-PAGE gel of the MT pelleting assay shown in Fig. 2B. Two gels were loaded with different input quantities: 40 and 2.5% of total samples. The positions of tubulin and mCherry-MAP7-Ct truncation on the gel are indicated on the right. (H) Images showing in vitro polymerized MTs, with HiLyte 488-labeled tubulin, Rhodamine-labeled MT seeds and mCherry-MAP7-Ct. Images were obtained on a TIRF microscope. Rhodamine labeled GMPCPP-stabilized MT seeds are indicated (red arrow). The mCherry-MAP7-Ct image (also showing MT seeds) is shown on the right with linearly increased brightness/contrast (ImageJ software).

**Figure S2.**
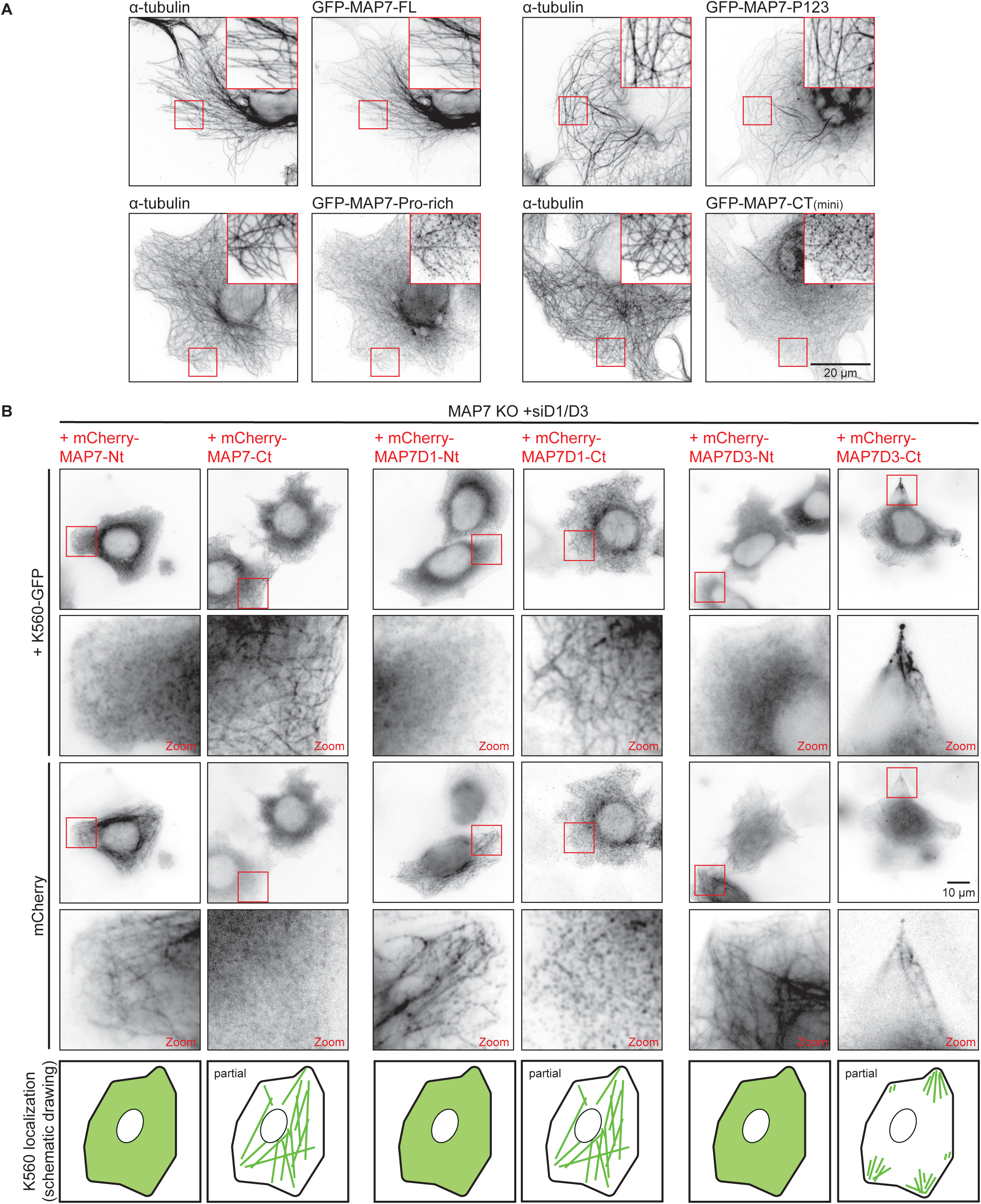
Kinesin-1 recruitment to MTs by the C-termini of MAP7 family proteins. (A) COS7 cells overexpressing indicated GFP-tagged MAP7 constructs co-stained for α-tubulin. Zooms are indicated with red boxes. (B) Wide field images of MAP7 KO + siMAP7D1/D3 HeLa cells overexpressing K560-GFP with mCherry-tagged MAP7-Nt and -Ct constructs. Enlargements of images indicated with a red squared box are shown in the panel row below. A schematic and representative drawing of K560-GFP localization for each condition is shown at the bottom.

**Figure S3.**
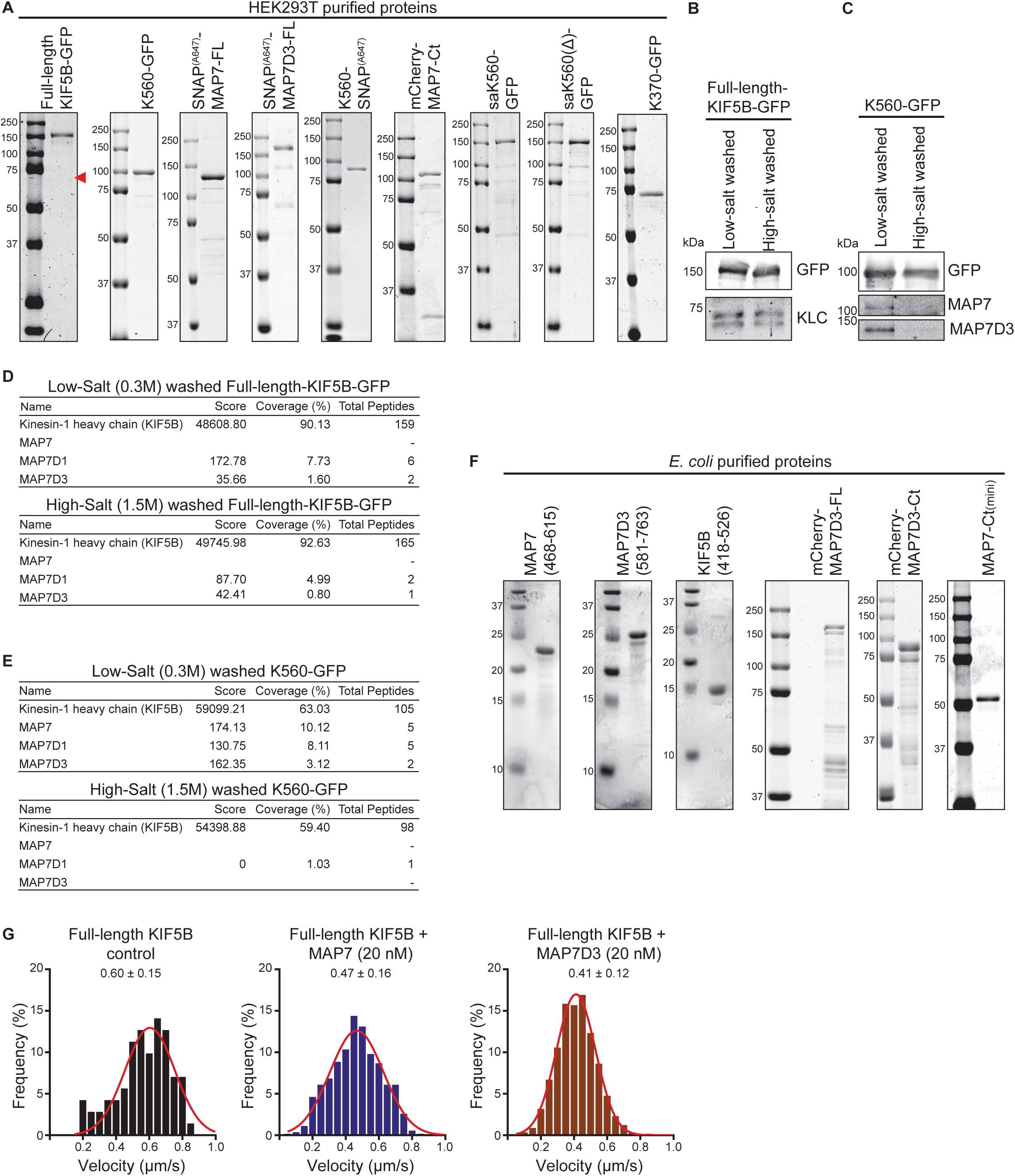
Overview and analysis of purified proteins. (A) Proteins purified from HEK293T cells used in this study analyzed by SDS-PAGE. (B, C) Western blot analysis of purified kinesins washed with low (0.3 M) or high-salt (1.5 M NaCl). Antibodies against GFP, KLC, MAP7 and MAP7D3 were used, GFP serves as a loading control for both experiments. (D, E) Mass-Spectrometry analysis of purified kinesins washed with low (0.3 M) or high-salt (1.5 M NaCl) buffer. (F) Proteins purified from *E. coli* used in this study analyzed by SDS-PAGE. (G) Histograms of full-length kinesin-1 velocities in control conditions or in the presence of 20 nM MAP7 or MAP7D3. Red lines show fitting with Gaussian distributions, mean values ±SD are indicated in the plot; n = 71 (control), n = 542 (MAP7), n = 568 (MAP7D3), from two or three independent experiments.

**Figure S4.**
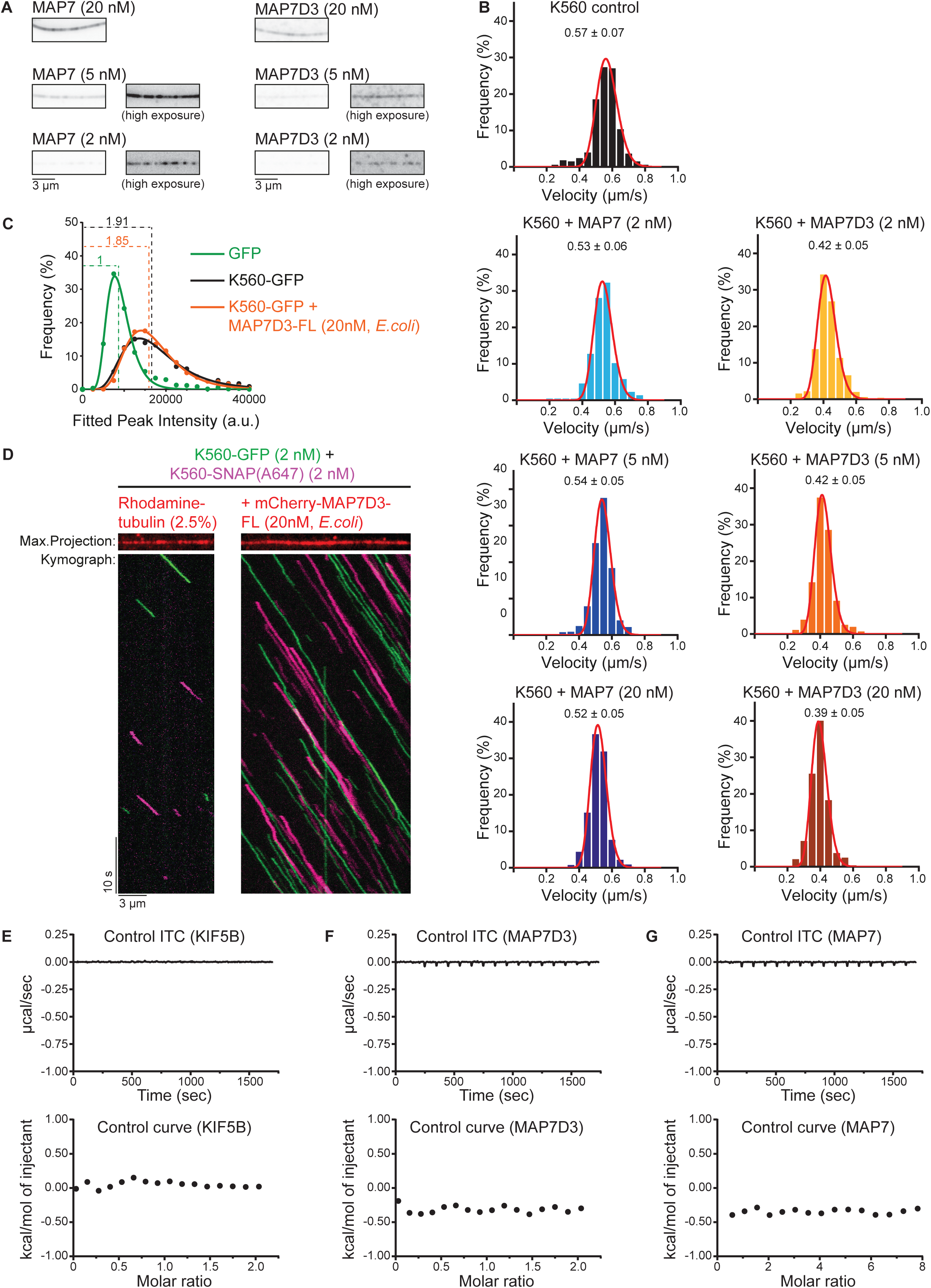
Velocities and oligomerization state of kinesin constructs in different conditions. (A) Images showing increasing concentrations of SNAP(Alexa647)-tagged MAP7 or MAP7D3 on dynamic MTs in vitro. Images were obtained with identical laser power and exposure time on a TIRF microscope. Panels on the right show replicate images from the left with linearly increased brightness/contrast (ImageJ software). (B) Histograms of K560-GFP velocities in control conditions or in the presence of indicated proteins. Red lines show fitting with Gaussian distributions, mean values with standard deviation are indicated in the plot, n = 241 (control), n = 351 (MAP7, 2 nM), n = 614 (MAP7, 5 nM), n = 361 (MAP7, 20 nM), n =257 (MAP7D3, 2nM), n = 436 (MAP7D3, 5 nM) and n = 303 (MAP7D3, 20 nM) from two independent experiments. (C) Histograms of fluorescence intensities of single GFP molecules (immobilized on coverslips) and K560-GFP moving on MTs with or without mCherry-MAP7D3-FL (purified from *E. coli*) in two separate chambers on the same coverslip (dots) and the corresponding fits with lognormal distributions (lines). n = 858 (GFP), n = 1640 (K560-GFP) and n = 4137 molecules (K560-GFP + mCherry-MAP7D3-FL); fluorophore density was approximately 0.01 µm^−2^ for GFP and K560-GFP proteins were analyzed from 2-10 MTs per movie. Dashed lines show corresponding relative median values. (D) Representative kymographs of 1:1 mixed K560-GFP (green) and K560-SNAP(Alexa647) (magenta) moving on dynamic MTs with or without mCherry-MAP7D3-FL (purified from *E.coli*). Maximum intensity projections show rhodamine-labeled MTs (control) or mCherry-MAP7D3-FL labeled MTs in red. (E-G) Control ITC experiments of KIF5B, MAP7D3 and MAP7. Top panels: Enthalpograms of the respective titrations. Lower panel: Black dots represent the integrated heat change.

**Figure S5.**
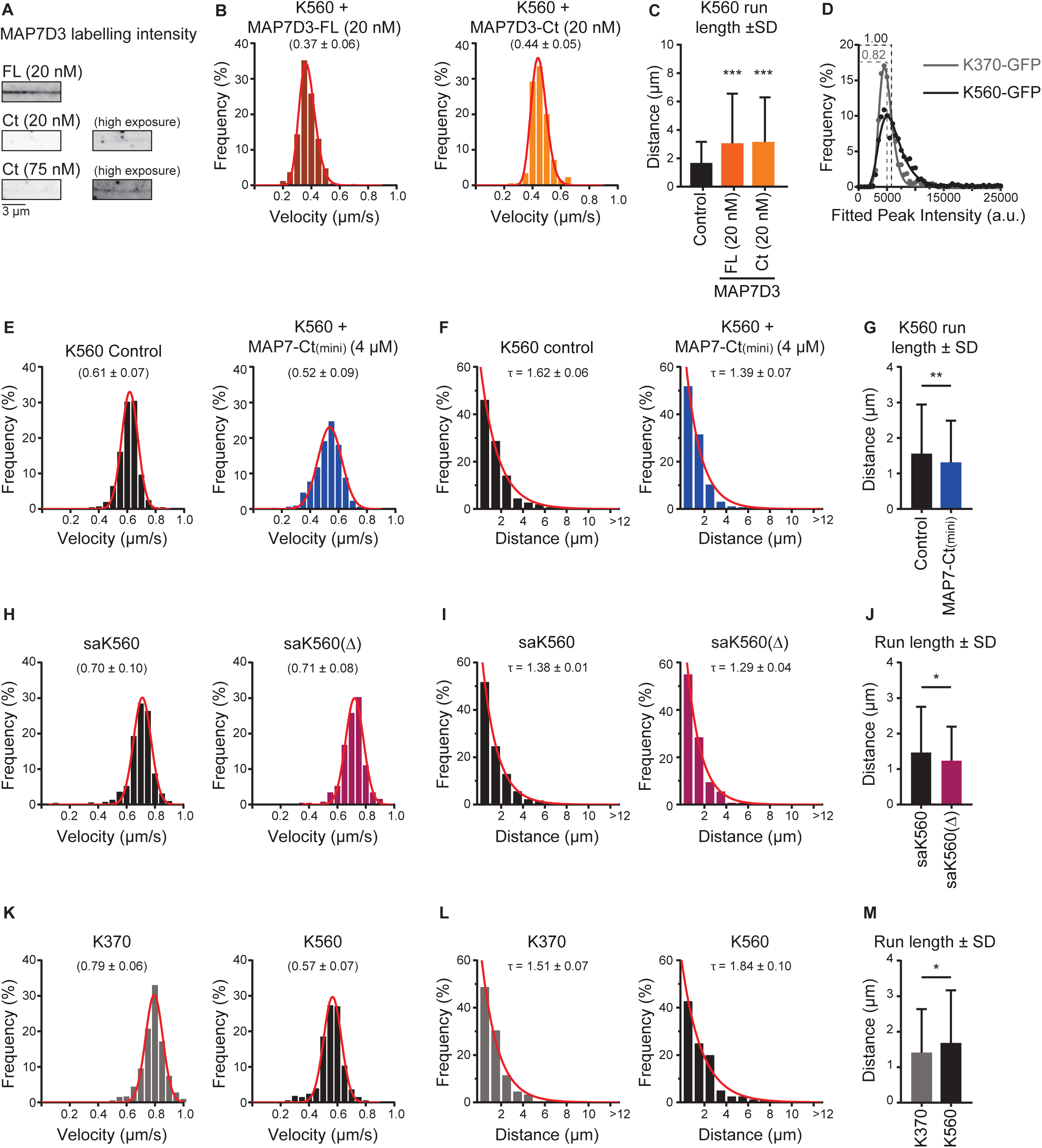
Kinesin motility parameters. (A) Images showing increasing concentration of mCherry-tagged MAP7D3 full-length or Ct (purified from *E.coli*) on dynamic MTs in vitro. Images were obtained with identical laser power and exposure time on a TIRF microscope. Panels on the right show replicate images from the left with linearly increased brightness/contrast (ImageJ software). (B) Histograms of K560-GFP velocities in the presence of indicated MAP7D3 proteins (purified from *E. coli*). Red lines show fitting with Gaussian distributions; mean values with standard deviation are indicated in the plot, n = 271 (MAP7D3-FL) and n = 209 (MAP7D3-Ct). (C) Quantification of K560-GFP run length in control condition or in the presence of the indicated MAP7D3 proteins (purified from *E.coli*). *** p<0.001, Mann-Whitney U test. n = 241 (control), n = 271 (MAP7D3-FL, 20 nM), n = 209 (MAP7D3-Ct, 20 nM). (D) Histograms of fluorescence intensities of K370-GFP and K560-GFP motors moving on MTs in two separate chambers on the same coverslip (dots) and the corresponding fits with lognormal distributions (lines). n = 639 (K370-GFP) and n = 1337 molecules (K560-GFP); motor proteins were analyzed from 2-10 MTs per movie. Dashed lines show corresponding relative median values. (E, H, K) Histograms of kinesin velocities. Red lines show fitting with Gaussian distributions; mean values with standard deviation are indicated in the plot, (E) n = 404 (control) and n = 648 (MAP7-Ct(mini)), (H) n = 804 (saK560) and n = 380 (saK560(∆)), (K) n = 723 (K370) and n = 241 (K560). (F, G, I, J, L, M) Kinesin run lengths were quantified and are shown as a histogram distribution with a fitted exponential decay curve (red), with indicated rate constants (tau) as a measure of mean run length (F, I, L) or with a bar graph (G, J, M) ** p <0.01, * p <0.05, Mann-Whitney U test. n numbers correspond to those of the preceding panels showing kinesin velocities.

## Videos

**Video 1. Imaging of light-induced nuclear export of K560-LEXY with GFP-MAP7**

Sequential dual-color video of GFP-MAP7 (left) and K560-mCherry-LEXY (right) in KIF5B KO HeLa cells. The video was acquired at 5 sec per frame over the course of 4 min on a spinning disc confocal microscope setup. Video corresponds to Fig. 4D.

**Video 2. Imaging of light-induced nuclear export of K560-LEXY with GFP-MAP7D3**

Sequential dual-color video of GFP-MAP7D3 (left) and K560-mCherry-LEXY (right) in KIF5B KO HeLa cells. The video was acquired at 5 sec per frame over the course of 4 min on a spinning disc confocal microscope setup. Video corresponds to Fig. 4G.

## Notes

#### Summary of Updates

The revised paper includes data on the regulation of full length kinesin-1 by MAP7 family members in vitro, on the role of unstructured linker sequences of MAP7 in microtubule binding and additional data supporting allosteric regulation of kinesin-1 by MAP7 proteins.

